# AAV-mediated gene therapy for Sialidosis

**DOI:** 10.1101/2023.11.10.566667

**Authors:** Diantha van de Vlekkert, Huimin Hu, Leigh E. Fremuth, Scott A. Brown, Jason A. Weesner, Elida Gomero, Yvan Campos, Alessandra d’Azzo

## Abstract

Sialidosis is a glycoprotein storage disease caused by deficiency of the lysosomal sialidase NEU1, which leads to pathogenic accumulation of sialylated glycoproteins and oligosaccharides in tissues and body fluids. The disease belongs to the group of orphan disorders with no therapy currently available. Here, we have tested the therapeutic potential of AAV-mediated gene therapy for the treatment of sialidosis in a mouse model of the disease. One-month-old *Neu1^−/−^* mice were co-injected with two scAAV2/8 vectors, expressing NEU1 and its chaperone PPCA, and sacrificed at 3 months post-injection. Treated mice were phenotypically indistinguishable from their WT controls. Histopathologically, they showed diminished or absent vacuolization in cells of visceral organs, including the kidney, as well as the choroid plexus and other areas of the brain. This was accompanied by restoration of NEU1 activity in most tissues, reversal of sialyl-oligosacchariduria, and normalization of lysosomal exocytosis in the CSF and serum of treated mice. AAV injection prevented the occurrence of generalized fibrosis, which is a prominent contributor of disease pathogenesis in *Neu1^−/−^* mice and likely in patients. Overall, this therapeutic strategy holds promise for the treatment of sialidosis and may be applicable to adult forms of human idiopathic fibrosis with low NEU1 expression.

## Introduction

Neuraminidase 1 (NEU1) is essential for the hydrolysis of sialo-glycoconjugates in lysosomes, which the enzyme initiates by cleaving terminal sialic acid residues from their glycan chains. In mammalian tissues NEU1 is found in a high molecular weight complex with two other lysosomal enzymes, the serine carboxypeptidase protective protein/cathepsin A (PPCA) and the glycosidase, β-galactosidase (β-Gal). Interaction with PPCA is a prerequisite for the proper localization, stability and activation of NEU1. In absence of a functional PPCA, NEU1 is enzymatically silent ^1,2^.

Congenital deficiency of NEU1 in humans affects lysosomal catabolism of sialo-glycoconjugates, resulting in the lysosomal storage disease (LSD), sialidosis or mucolipidosis I ^3,4^. Patients with sialidosis are classified into two subtypes based on the age of onset and severity of their symptoms, as well as residual enzyme activity. Type I sialidosis is the attenuated form of the disease with onset during adolescence, while type II sialidosis is the severe, neuropathic form of the disease, which can be further classified into congenital or hydropic, infantile and juvenile ^4,5^. Type I patients have normal intellectual ability and are mostly asymptomatic during childhood or early adolescence. They are typically diagnosed in the second decade of life, when they develop symptoms, such as myoclonus, gait abnormalities and progressive visual loss, associated with bilateral macular cherry-red spots ^4,5^. Type II patients with the acute congenital form of the disease present with facial edema and inguinal hernias, and can develop hydrops fetalis, neonatal ascites, or both, leading to stillbirth or death shortly after birth. The infantile/juvenile type II patients have coarse facial features, hepatosplenomegaly, dysostosis multiplex, skeletal dysplasia, macular cherry-red spots, hearing loss, angiokeratoma, myoclonus, cardiomyopathy and mental retardation ^4,6^. Lysosomal accumulation of sialylated glycoproteins and oligosaccharides in tissues and body fluids (oligosacchariduria) are diagnostic of both disease subtypes.

*Neu1* knockout mice (*Neu1^−/−^*) closely resemble type II sialidosis ^7^. They present with signs of the disease at birth, associated with severe growth retardation and, in a percentage of newborns, death at weaning^7^. The disease progresses into a severe systemic and neurodegenerative condition, leading to premature death (4-6 months). During the course of their lifespan, affected mice show dysmorphic features, progressive edema and oligosacchariduria^7^. This sialidosis model was extensively used to study mechanisms of pathogenesis and to test preclinical therapeutic modalities. During these studies, Neu1 was identified as a central negative regulator of the ubiquitous process of lysosomal exocytosis that Neu1 restrains by desialylating the lysosomal associated membrane protein 1, Lamp1, extending its half-life ^8–10^. Loss of Neu1 activity leads to an increased number of Lamp1+ lysosomes docked at the plasma membrane (PM) poised to fuse with the PM and release their content extracellularly. Exacerbation of lysosomal exocytosis has since been established as a central pathogenic mechanism in the *Neu1^−/−^* mouse model, that likely reflect known clinical manifestations of sialidosis patients ^10–15^. For instance, excessive lysosomal exocytosis in the *Neu1^−/−^* connective tissue has been linked to the development of muscle fibrosis responsible for muscle fiber degeneration and muscle atrophy ^13,15^. A similar paradigm applied to the brain of *Neu1^−/−^* mice was shown to contribute to the progressive deposition of β−amyloid resembling Alzheimer’s disease pathology and leading to neurodegeneration ^11^.

Currently, there is no target treatment or cure available for type I or II sialidosis. Patients receive supportive care and symptomatic relief, primarily for the management of myoclonus and seizures^4,5^. We have previously tested different therapeutic approaches in various Neu1 deficient mouse models, including enzyme replacement therapy (ERT)^16^, chaperone-mediated gene therapy with PPCA^17^ and pharmacological and dietary compounds^18^. Specifically, treatment of a type 1 sialidosis model with a single dose of a liver specific AAV vector expressing PPCA enhanced residual Neu1 activity and prevented kidney pathology and oligosacchariduria^17^.

These encouraging results have set the stage for the current preclinical study exploring the combined use of AAV vectors expressing NEU1 and PPCA for the treatment of sialidosis. A cohort of *Neu1^−/−^* mice were injected with two self-complementary recombinant AAV2/8 (scAAV) vectors expressing *NEU1* and *CTSA* under the control of a constitutive cytomegalovirus (CMV) early enhancer promotor. Treatment prevented disease progression, ameliorated the neuropathological signs and the occurrence of fibrosis, two difficult-to-treat phenotypes in sialidosis patients.

## Results

### Biodistribution of AAV vectors and immune response in injected mice reflect positive therapeutic outcome

To investigate whether the administration of a single combined dose of scAAV2/8-CMV-*NEU1* (2×10^12^ vg/mouse) and scAAV2/8-CMV-*CTSA* (1×10^12^ vg/mouse) (scAAV-N/P) could revert or prevent the development of typical pathological manifestations in the sialidosis mice, we co-injected a cohort of 1-month-old *Neu1^−/−^* mice that were sacrificed at 3 months post injection. Treated mice appeared almost indistinguishable from their control littermates: their dysmorphic facial features and progressive edema of the eyelids and paws, two prominent phenotypes of the disease, were largely resolved (Figure 1A).

**Figure 1.**
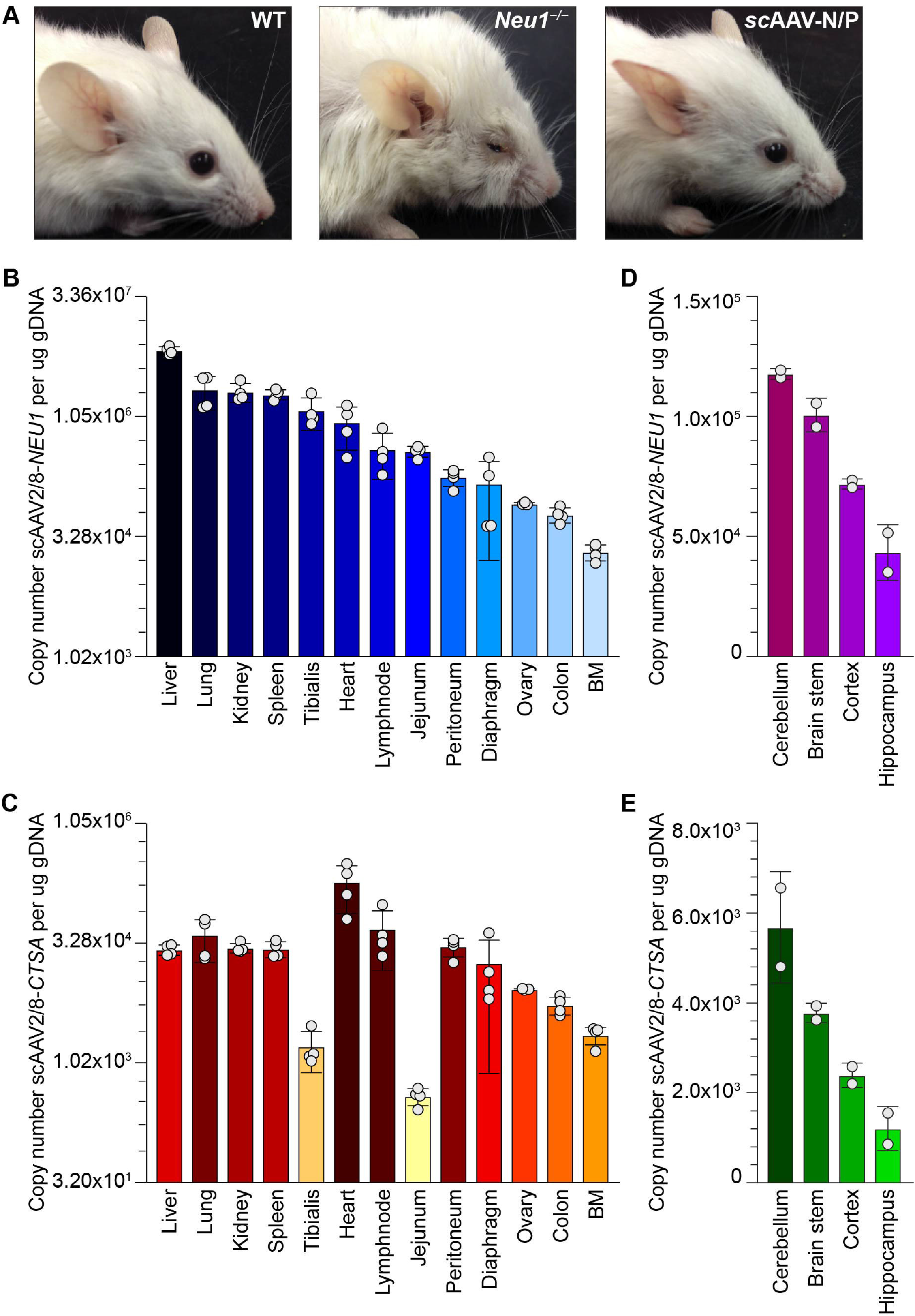
Gross appearance of *Neu1^−/−^* treated mice and tissue biodistribution of recombinant AAV vectors. (A) Representative images of edema of the eyelids and disheveled coat of *Neu1^−/−^* mice, features that were corrected in scAAV-N/P treated mice. (B and C) Copy number of scAAV2/8-CMV-*NEU1* (B) and scAAV2/8-CMV-*CTSA* (C) in multiple organs of injected mice at 3 months post injection, measured per μg of genomic DNA. *n* = 4; ovary: *n* = 3. (D and E) scAAV2/8-CMV-*NEU1* (D) and scAAV2/8-CMV-*CTSA* (E) copy number in cerebellum, brain stem, cortex and hippocampus. *n* = 2.

We then assessed the vectors’ biodistribution by measuring the copy number of recombinant scAAV2/8-CMV-*NEU1* and scAAV2/8-CMV-*CTSA* in the total genomic DNA extracted from multiple organs of treated mice. We found that the highest *NEU1* viral load (average of 7.1×10^6^) was detected in the liver, followed by the lung, kidney, spleen and tibialis anterior (Figure 1B). The other tissues had 1 to 2 log_2_ lower amounts of viral genome compared to the liver, with the BM containing the lowest copy number (average 2.09×10^4^) (Figure 1B). Testing the viral load of *CTSA* in the same tissues showed a different biodistribution pattern compared to *NEU1* with the highest viral load in the heart (average 1.93×10^5^), followed by the lymph nodes, lung, peritoneum, kidney, spleen, liver and diaphragm (Figure 1C). The remaining tissues had a 2 log_2_ lower amount of viral genome compared to the heart, with the lowest amount found in the jejunum (3.9×10^2^) (Figure 1C). The average viral loads for both vectors in different regions of the brain, namely the cerebellum, brain stem, cortex and hippocampus, were remarkably high (Figures 1D and 1E), indicating that the virus had crossed the blood brain barrier. We also tested whether treated mice would mount an adaptive immune response toward the vector (AAV2/8) and/or the transgene product (NEU1 and PPCA). For this purpose, we tested serum samples from injected mice and controls at 4 months of age using ELISA. No antibodies specific for either human NEU1 or PPCA were detected in any tested samples (Figures S1A and S1B). However, we did measure antibodies against the viral capsid in the injected samples (Figures S1C and S1D), which were likely due to the use of a self-complementary double-stranded AAV vector, rather than a single-stranded vector, as reported previously^19^.

### Expression of therapeutic NEU1 and PPCA is detected in different brain regions

It is well established that the brain is an exceedingly difficult tissue to treat via gene therapy due to the presence of the blood-brain-barrier, especially following a systemic injection route ^20^. Thus, we wanted to determine whether the expression of the transgene products in different brain regions of treated mice would parallel the biodistribution of the two AAV vectors. Both NEU1 and PPCA proteins were readily detectable by immunohistochemistry (IHC) in the epithelial cells of the choroid plexus (Figures 2A and 2B), but only in sparse neurons of the CA3 region of the hippocampus and in a few Purkinje cells of the cerebellum (Figures S2A – S2D). The efficacy of infection was further tested by measuring the enzyme activities of NEU1 and PPCA in the same brain regions of scAAV-N/P injected mice. Slightly increased NEU1 activity was measured in the brain stem, cortex and cerebellum of the treated mice compared to the *Neu1^−/−^* non-injected controls, although the values did not reach statistical significance (Figure 2C). Cathepsin A (CA) activity instead followed a different pattern. As we have previously established in sialidosis patients’ fibroblasts, deficiency of NEU1 is associated with a variable increase in the CA activity^18^. We now confirmed this paradigm in the *Neu1^−/−^* mice, where significantly increased activities for both CA and β-Gal, the third member of the lysosomal multienzyme complex, were measured in the different regions of the *Neu1^−/−^* brain, highest in the hippocampus, compared to the WT controls (Figures 2D and 2E). Interestingly, the hippocampus of the injected mice was the best responding region of the brain, showing a significant decrease (35%) in CA activity towards its normal levels, whereas the other 3 regions were unchanged (Figure 2D). Instead, β-Gal activity was reduced in all four brain regions of the injected mice (Figure 2E), confirming the dependency of β-Gal on the localization and activity levels of NEU1 ^21^. Two other lysosomal enzymes, β-hexosaminidase (β-Hex) and α-mannosidase (α-Man), known to be influenced by the deficiency of Neu1, were also used as proxy method to assess efficacy of treatment. These enzyme activities were significantly increased in all brain regions of the *Neu1^−/−^* mice compared to the WT controls (Figures 2F and 2G), but their values were again largely reduced in the same regions of the injected mice (Figures 2F and 2G). The data suggest that only minuscule increase in NEU1 and CA activities in the brain of scAAV-N/P injected mice was sufficient to partially normalize the levels of the other lysosomal enzymes.

**Figure 2.**
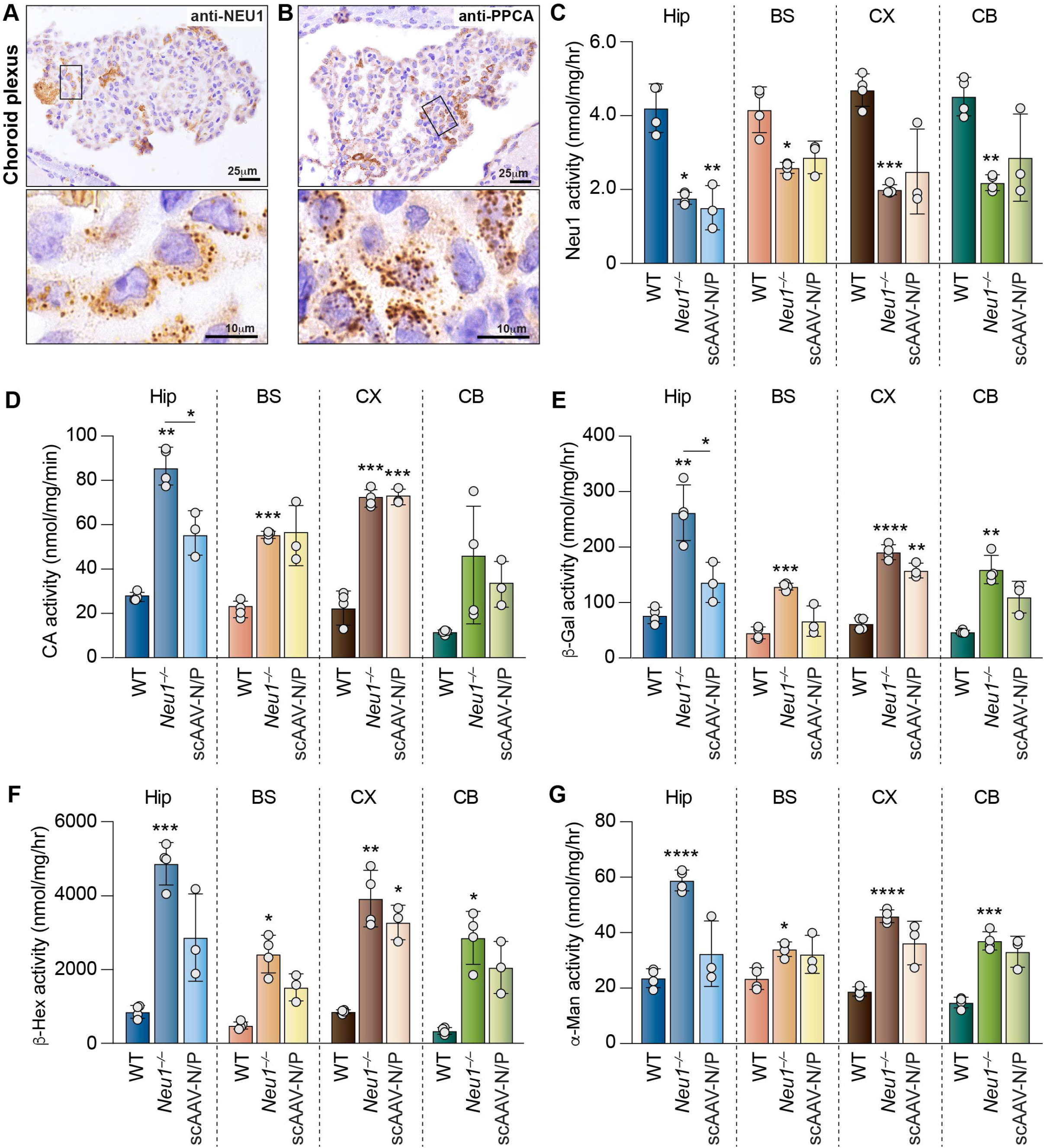
Partially restored expression of NEU1 and PPCA in different brain regions of *Neu1^−/−^*mice treated with scAAV-N/P normalizes the levels of other lysosomal enzymes. (A – D) IHC of NEU1 and PPCA in the CP epithelial cells of treated mice. Scale bar: 25 μm. Lower panels are zoomed in images of boxed areas. Scale bar: 10 μm. Brown puncta depict compartmentalized/lysosomal NEU1 and PPCA. (C – G) Enzyme activity levels for NEU1 (C), CA (D), β-Gal (E), β-Hex (F) and α-Man (G) in the hippocampus (Hip), brain stem (BS), cortex (CX) and cerebellum (CB) of WT (*n* = 4), *Neu1^−/−^*(*n* = 4) and scAAV-N/P treated (*n* = 3) mice. Values are expressed as mean ± SD. Statistical analyses were performed using One-way ANOVA; \**p* < 0.05, \*\**p* < 0.01, \*\*\**p* < 0.001, \*\*\*\**p* < 0.0001.

### Reversal of histopathology and reduction of microgliosis occur in the brain of scAAV-treated Neu1^−/−^ mice

Epithelial/endothelial cells and macrophages, including microglia, are primary targets of pathogenesis in sialidosis ^4,10,11^. The epithelial cells of the choroid plexus (CP) follow this pattern, showing signs of the disease, i.e., numerous storage-filled lysosomes, early in the life of *Neu1^−/−^* mice (Figure S2E). After scAAV-N/P treatment, these cells were fully corrected, as seen on H&E stained brain sections, and the CP of treated mice appeared morphologically indistinguishable from that of the WT mice (Figure S2E). In addition, IHC with Lamp1 antibody revealed normalization of the levels of Lamp1, in different brain regions of the treated mice, including the CP epithelium (Figure 3A), the CA3 region of the hippocampus and scattered glial cells in the cerebellum (Figure S3A and S3B). This finding was particularly relevant because we have shown previously that Lamp1 accumulation downstream of Neu1 deficiency is paralleled by excessive lysosomal exocytosis. To measure the extent of lysosomal exocytosis in treated and untreated mice we assayed the activity of β-Hex in their cerebrospinal fluids (CSF) and sera. We found that the increased β-Hex activity in both samples from the *Neu1^−/−^* mice was significantly decreased after treatment, confirming that reduced levels of Lamp1 in treated mice were accompanied by decreased lysosomal exocytosis (Figures 3B and 3C). However, correction of this phenotype was not sufficient to prevent the formation of amyloid deposits in scAAV-N/P injected mice, likely due to the limited NEU1 expression and activity in neurons (Figure S3C).

**Figure 3.**
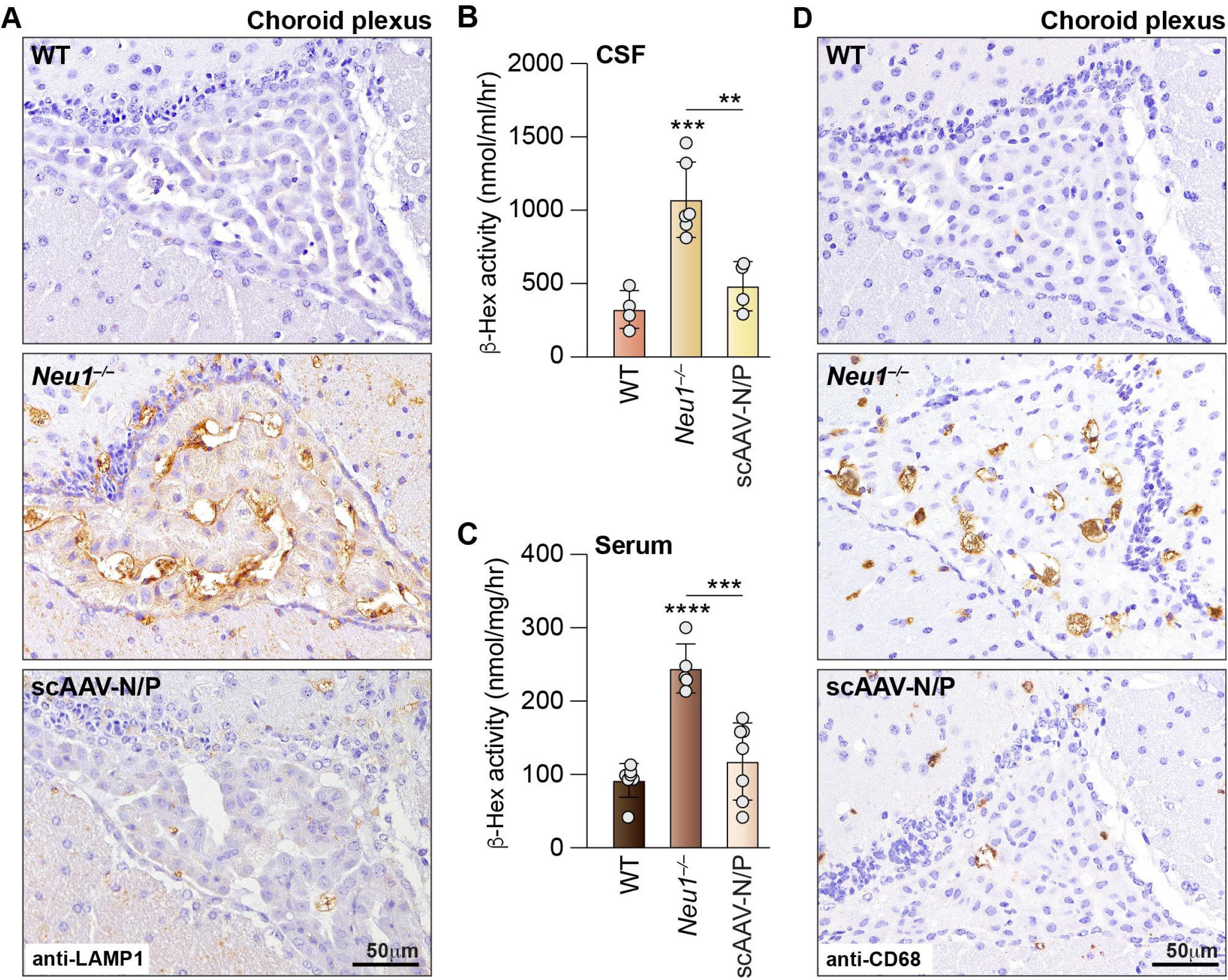
scAAV-N/P treatment reduces Lamp1 levels, lysosomal exocytosis in the CSF and serum, and brain inflammatory marker, CD68. (A) Representative IHC images of the choroid plexus from WT, *Neu1^−/−^*and scAAV-N/P treated mice, using anti-Lamp1 antibody. Scale bar: 50 µm. (B) β-Hex enzyme activity measured in the CSF from WT (*n* = 4), *Neu1^−/−^* (*n* = 6) and scAAV-N/P treated (*n* = 4) mice. (C) β-Hex enzyme activity measured in sera from WT (*n* = 7), *Neu1^−/−^* (*n* = 5) and scAAV-N/P treated (*n* = 7) mice. Values are expressed as mean ± SD. Statistical analyses were performed using One-way ANOVA; \*\**p* < 0.01, \*\*\**p* < 0.001, \*\*\*\**p* < 0.0001. (D) Representative IHC images of the choroid plexus from WT, *Neu1^−/−^*and scAAV-N/P treated mice using anti-CD68 antibody. Scale bar: 50 µm.

Another suitable parameter to determine therapeutic efficacy in scAAV-N/P treated *Neu1^−/−^* mice is microglia activation. We have previously established that tissue macrophages and microglia are one of the most affected cell types in *Neu1^−/−^* mice ^7^. To assess the status of microglia activation before and after AAV injection we performed IHC of brain sections from WT, *Neu1^−/−^* and scAAV-N/P mice with anti-CD68 antibody. CD68 is a widely used, specific marker of pro-inflammatory, activated monocyte cells, which include microglia. The *Neu1^−/−^* mice showed clearly increased CD68 staining of microglia, particularly in the CP, the CA3 region of the hippocampus and the cerebellum compared to the WT controls, which had no detectable CD68+ cells (Figures 3D, S3C and S3D). Mice treated with scAAV-N/P showed diminished CD68 staining at 3 months post injection (Figures 3D, S3C and S3D), indicating a decrease in microglia-mediated neuroinflammation.

These combined results showed that a single IV co-injection of scAAV2/8-CMV-*NEU1* and scAAV2/8-CMV-*CTSA* promotes the expression of both therapeutic proteins resulting in partial correction of different brain areas.

### Restored NEU1 activity in treated Neu1^−/−^ mice reverse fibrosis in skeletal muscle and heart

Extensive and progressive expansion of the connective tissue in *Neu1^−/−^* skeletal and heart muscles is the cause of muscle fiber degeneration, muscle atrophy ^13,15^, and heart fibrosis. This phenotype has been linked to the effects of Neu1 deficiency on fibroblasts that retain a partially differentiated and activated state, characterized by increased Lamp1 levels, and excessive exocytosis of lysosomal contents and exosome that propagate fibrotic signals to unaffected areas and bolster disease progression ^13^. For the evaluation of the muscle phenotype before and after treatment, we first tested the expression of NEU1 and PPCA proteins in skeletal muscle and heart by IHC. Sustained expression of both proteins was detected in cardiomyocytes throughout the heart of treated mice (Figures 4A, S4A and S4B). Instead in the gastrocnemius (GA) muscle positive staining for PPCA was seen in virtually all fibers, while NEU1 expression appeared to be more intense in some fibers than others (Figure 4B and S4C and S4D).

**Figure 4.**
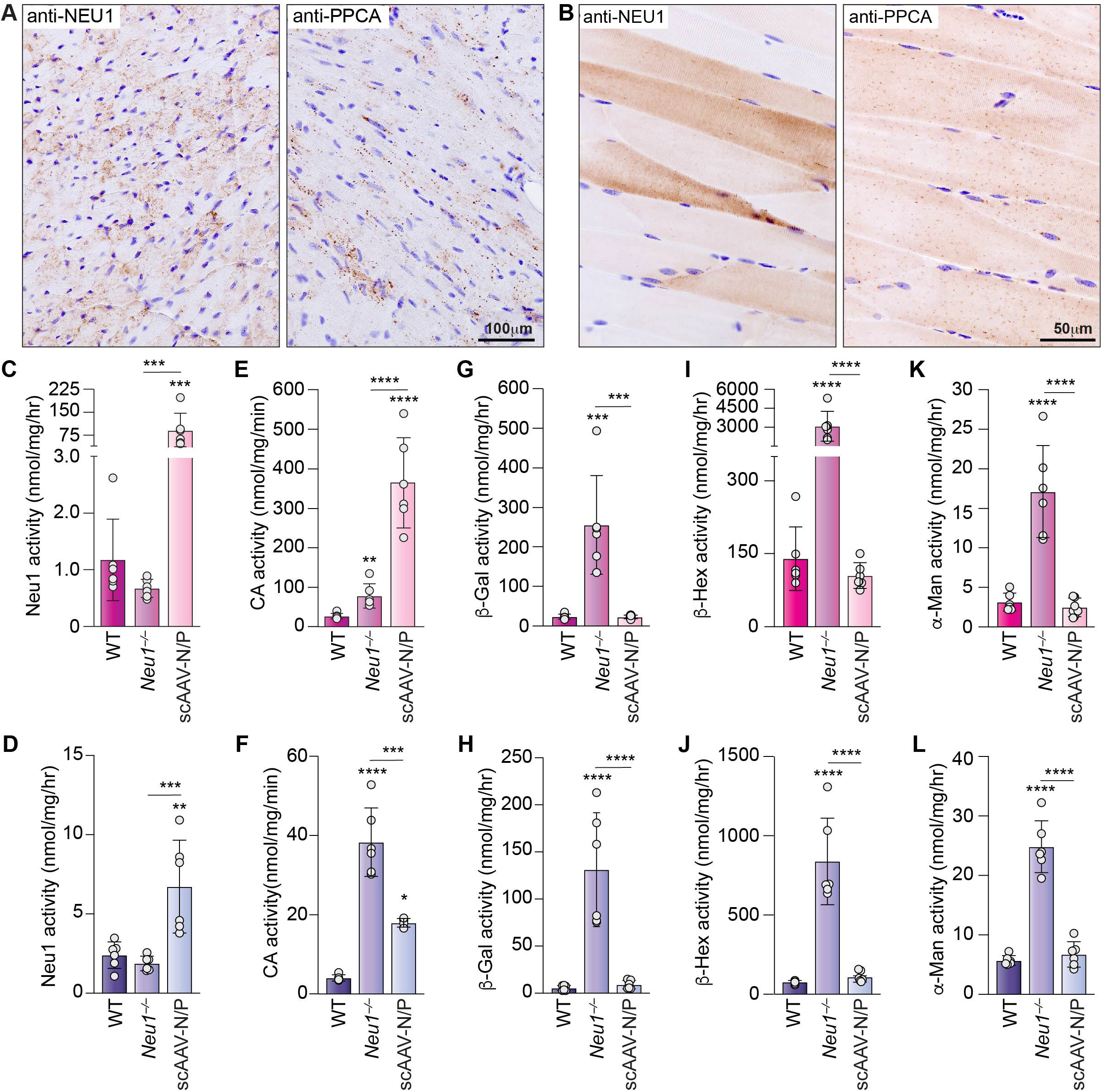
NEU1 and PPCA protein expression in heart and skeletal muscle corrects the activity of other lysosomal enzymes in *Neu1^−/−^* mice treated with scAAV-N/P. (A and B) Representative IHC images of heart (A) and GA skeletal muscle (longitudinal) (B) tissue sections from scAAV-N/P treated mice using anti-human NEU1 (left; A and B) and anti-human PPCA (right; A and B) antibodies. Scale bar: 100 µm and 50 µm. Brown puncta depict compartmentalized/lysosomal NEU1 and PPCA. (C – L) Enzyme activity levels measured in the heart and GA skeletal muscle from WT, *Neu1^−/−^* and scAAV-N/P treated mice: NEU1 (C and D), CA (E and F), β-Gal (G and H), β-Hex (I and J) and α-Man (K and L); *n* = 6. Values are expressed as mean ± SD. Statistical analysis was performed using the One-Way ANOVA; \*\**p* < 0.01, \*\*\**p* < 0.001, \*\*\*\**p* < 0.0001.

Efficacy of AAV infection was further determined by assaying the enzyme activities of NEU1 and PPCA in the heart and GA muscle of scAAV-N/P injected mice. Very high NEU1 and CA activities were measured in the heart tissue of treated mice (Figures 4C and 4D), likely due to the high transduction efficacy of cardiomyocytes ^22,23^. In the GA muscle of treated mice NEU1 activity was lower than in the heart but also robustly increased compared to untreated mice and WT controls (Figure 4E). Again, the increased CA activity in untreated *Neu1^−/−^* mice was reduced in the injected mice (Figure 4F). Also, in these tissues of the *Neu1^−/−^* mice, β-Gal activity was significantly higher (∼10x and ∼25x, respectively) compared to the WT controls, but the enzyme’s levels were normalized after treatment (Figures 4G and 4H). Similar to β-Gal, the increased activities ofβ-Hex and α-Man in *Neu1^−/−^* heart and GA muscle were restored to WT levels in treated mice (Figures 4I – L).

Correction of Neu1 activity was accompanied by clear improvement of the histopathology in heart and GA muscle of treated *Neu1^−/−^* mice. We paid particular attention to the connective tissue component ^13,15^. Extensive vacuolization of fibroblasts at the attachment site of the heart valves, including the mitral valve, was prominent in the *Neu1^−/−^* heart (Figure S5A). This structure was fully corrected in the treated mice and appeared morphologically comparable to that of the WT controls (Figure S5A). Similarly, the occurrence of fibrosis, characteristic of the *Neu1^−/−^* skeletal muscle ^13,15^ was prevented in treated mice (Figure S4B).

Normalization of tissue morphology at 3-months post injection (Figures S4A and S4B), was confirmed by staining heart and GA muscle sections from *Neu1^−/−^* mice and scAAV-N/P-treated mice with Masson’s Trichrome. The heart and GA muscle tissues from the treated mice showed no signs of increased collagen deposition that was evident in both tissues of the untreated *Neu1^−/−^* mice (Figures S4C and S4D). To further assess the degree of correction of the heart and muscle phenotype, we immune-stained sections of the heart and GA muscle with anti-Lamp1 and anti-CD68 antibodies. Lamp1 expression levels in both tissues of the treated mice were comparable to those in the control group (Figures 5A, 5B and S4E), and the number of CD68+ macrophages were also reduced (Figures 5C and 5D).

**Figure 5.**
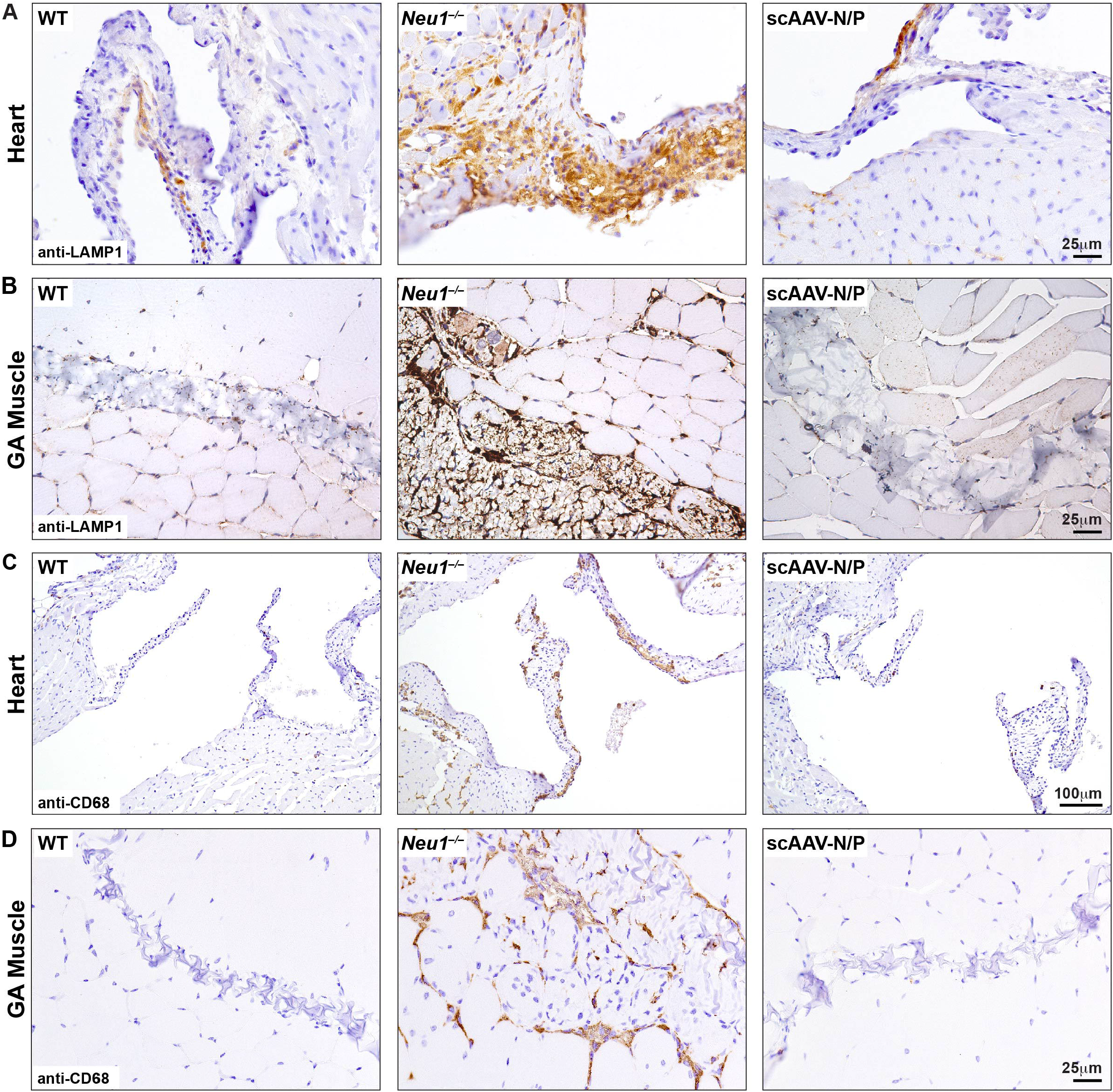
Reduced Lamp1 and CD68 levels in scAAV-N/P treated mice. (A and B) Normalization of Lamp1 staining in the heart (A) and skeletal muscle (B). Scale bar: 25 µm. (C and D) Reduction of the inflammatory marker CD68 in both muscle tissues; heart (C) and skeletal muscle (D). Scale bars: 100 µm and 25 µm.

### Partial correction of kidney morphology in treated mice is accompanied by reversal of sialyl-oligosacchariduria

We next examined the effects of scAAV-N/P treatment in the kidney, a difficult to treat organ and one of the primary target of sialidosis in children ^4^. Positive antibody staining of NEU1 and PPCA was detected in the proximal tubules and to some extent in the glomeruli of scAAV-N/P-injected mice (Figures 6A and 6B). We also measured a 2-fold increase in NEU1 activity in kidney tissue lysates from treated mice compared to *Neu1^−/−^* samples (Figure 6C). CA activity levels were slightly increased in the *Neu1^−/−^* kidney but were similar to WT levels after treatment (Figure 6D). Similarly, β-Gal activity, which was significantly higher (∼3x) than the WT controls in the untreated mice, was normalized in the injected mice (Figure 6E). Normalization of enzyme activities was also observed for β-Hex and α-Man, a finding consistent with the results obtained in other tissues (Figure 6F and 6G).

**Figure 6.**
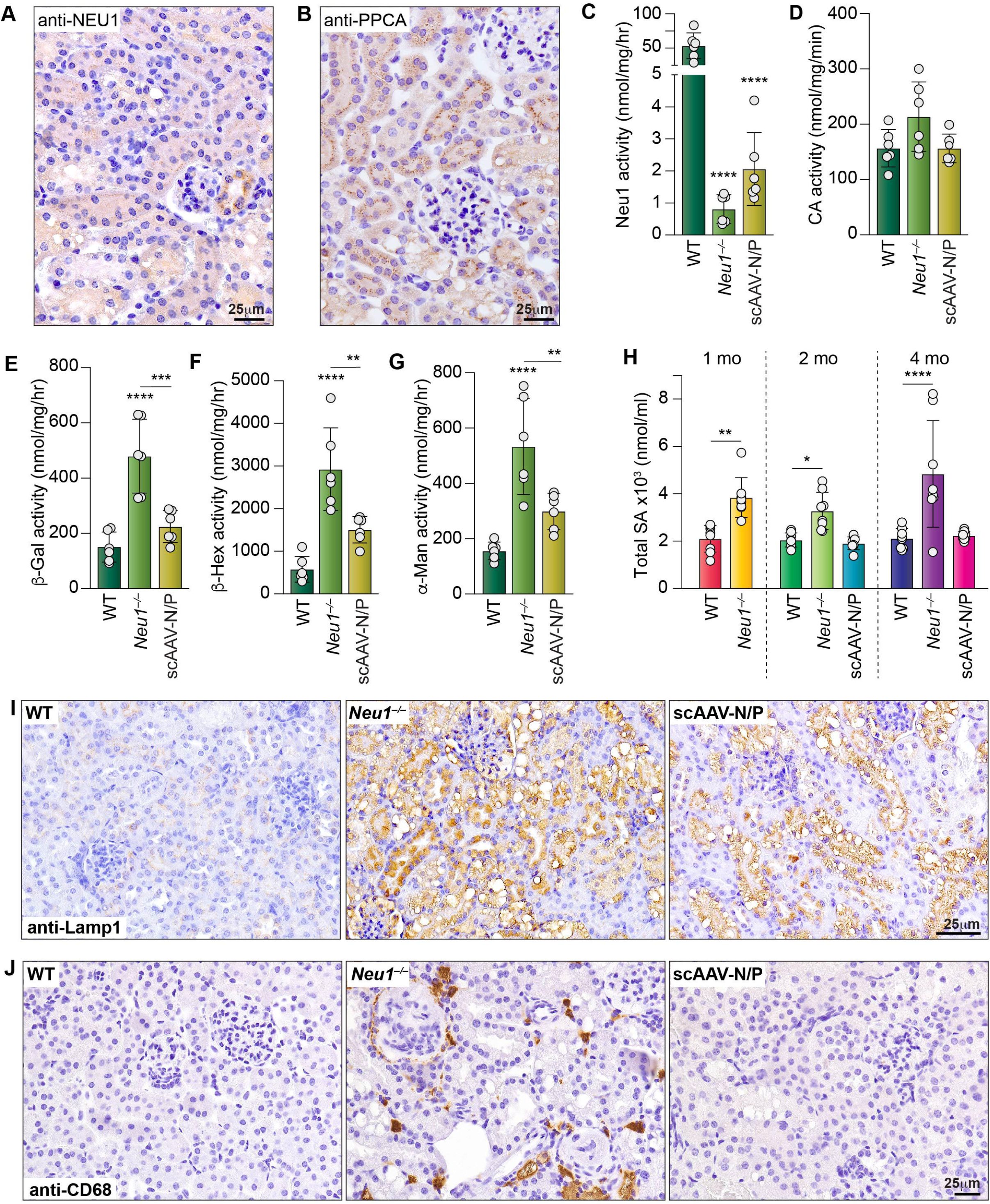
Expression of therapeutic NEU1 and PPCA in the kidney of treated mice normalizes other enzyme activities, resolves sialyl-oligosacchariduria and reduces Lamp1 and CD68 levels. (A and B) Representative IHC images of the kidney from scAAV-N/P treated mice using anti-human NEU1 (A) and anti-human PPCA (B) antibodies. Brown puncta depict compartmentalized/lysosomal NEU1 and PPCA. Scale bar: 25 µm. (C – G) Enzyme activity levels measured in the kidneys of WT, *Neu1^−/−^* and scAAV-N/P treated mice: NEU1 (C), CA (D), β-Gal (E), β-Hex (F) and α-Man (G), (*n* = 6). Values are expressed as mean ± SD. Statistical analysis was performed using the One-Way ANOVA; \*\**p* < 0.01, \*\*\**p* < 0.001, \*\*\*\**p* < 0.0001. (H) Total sialic acid levels measured in the urine of WT, *Neu1^−/−^* and *Neu1^−/−^*-scAAV-N/P mice at 1 month, 2 months, and 4 months of age (*n* = 8). Values are expressed as mean ± SD. Statistical analysis was performed using the One-Way ANOVA; \**p* < 0.05, \*\**p* < 0.01, \*\*\*\**p* < 0.0001. (I and J) Representative IHC images of the kidney from WT, *Neu1^−/−^*and scAAV-N/P treated mice stained with anti-Lamp1 (I) and anti-CD68 (J) antibodies. Scale bar: 25 µm.

Deficiency of NEU1 causes the accumulation of sialic acid-containing glycoconjugates in tissues and sialyl-oligosacchariduria, features that can be used as diagnostic markers and as measure of disease severity^4,7^. Kidney histopathology of *Neu1^−/−^* mice at 1 month and 4 months of age showed progressive vacuolization of the proximal tubule epithelium and the glomeruli (Figures S6A and S6B). Treatment of the *Neu1^−/−^* mice with scAAV-N/P at 1 month of age prevented disease progression in the kidneys, showing reduced vacuolization of the tubular epithelium at 4 months of age compared to the untreated *Neu1^−/−^* mice (Figures S6A and S6B). We then tested if injection of scAAV-N/P resolved the sialyl-oligosacchariduria in urine samples collected from age-matched WT, *Neu1^−/−^* and injected mice. Increased levels of total sialic acids, indicative of sustained sialyl-oligosacchariduria, were measured in samples of *Neu1^−/−^* mice at all ages (Figure 6H). This characteristic phenotype was fully resolved in mice injected with scAAV-N/P (Figure 6H). Remarkably, treatment prevented the occurrence of sialyl-oligosacchariduria over time, albeit the tissue morphology of the kidney was not completely corrected.

We also stained kidney sections from WT, *Neu1^−/−^* and scAAV-N/P-treated mice for Lamp1. In the WT kidney, Lamp1 positive staining was only detected in the proximal tubules, whereas in the *Neu1^−/−^* tissue both the proximal tubule and the glomeruli showed increased Lamp1 staining (Figure 6I). Upon scAAV-N/P treatment, Lamp1 level was decreased by ∼50% (Figure 6I). Moreover, IHC of kidney sections from the same set of mice with anti CD68 antibody demonstrated that scAAV-N/P treatment resolved the inflammatory response. Contrary to the untreated mice that had increased number of CD68+ cells, especially surrounding the glomeruli, no CD68+ cells were detected in the kidney of treated mice, indicating reversal of the inflammatory response (Figure 6J).

### Full correction of histopathology in the liver and spleen of treated Neu1^−/−^ *mice*

Similar positive outcome post treatment was observed in the liver and spleen of scAAV-N/P injected mice compared to the untreated *Neu1^−/−^* mice. IHC of liver tissue from scAAV-N/P injected mice showed a very prominent NEU1 and PPCA staining in hepatocytes, Kupfer cells and endothelial cell of the sinusoids (Figure 7A and 7B). In the spleen, the highest number of NEU1 and PPCA positive splenocytes was evident in the red pulp (Figures 7C and 7D). In addition, treated mice showed no evidence of splenomegaly (Figures S6C and S6D), a feature of *Neu1^−/−^* mice that is linked to time-dependent extramedullary hematopoiesis ^10^. NEU1 activity in the liver of scAAV-N/P injected mice was 5x higher that in WT controls (Figure 7E), while in the spleen no significant changes in activity levels were measured after treatment (Figure 7F). CA activity, which was significantly increased in both liver and spleen of the *Neu1^−/−^* mice (Figures 7G and 7H), was normalized after treatment in the spleen (Figure 7H) but remained high in the liver (Figure 7G). This again could be due to the high viral load in the liver of treated mice and the sustained expression of the PPCA protein in hepatocytes. In addition, the activities of β-Gal, β-Hex and α-Man were restored to WT levels in both tissues of scAAV-N/P injected mice (Figures 7I – 7N).At the histopathology levels, H&E-stained sections of the *Neu1^−/−^* liver showed severe and widespread vacuolization of the hepatocytes, the epithelial cells (cholangiocytes) of the bile ducts, the endothelial cells of the sinusoids and the Kupffer cells (Figure S7A). Liver histopathology was fully corrected in treated mice (Figure S7A). Similarly, in the *Neu1^−/−^* spleen, the large areas of the red pulp showing extensive vacuolization of splenocytes appeared cleared of storage after treatment (Figure S7B).

**Figure 7.**
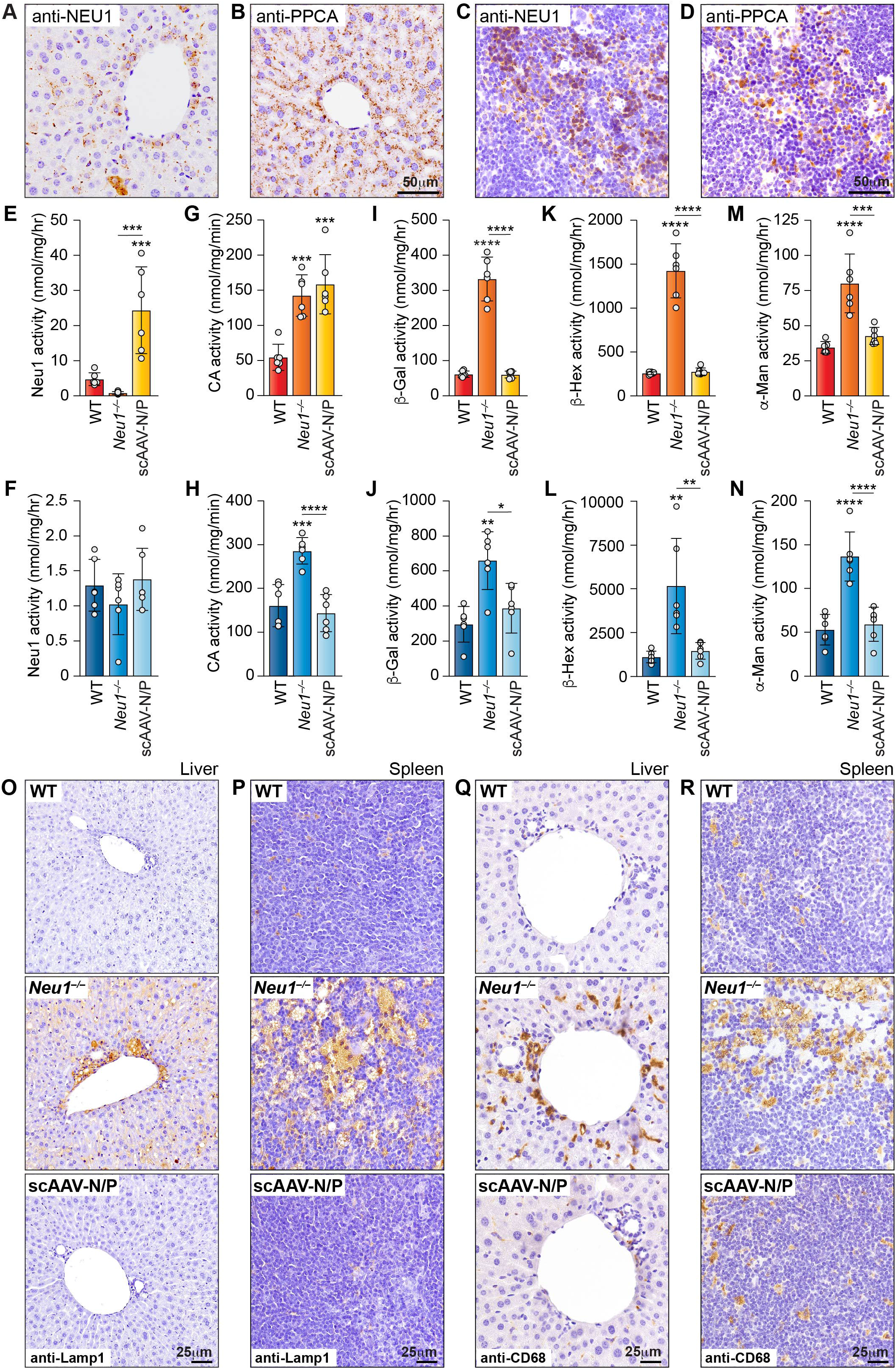
Expression of NEU1 and PPCA in liver and spleen of treated mice normalizes other enzyme activities and reduces Lamp1 and CD68 levels. (A – D) Representative IHC images of the liver (A and B) and spleen (C and D) from scAAV-N/P treated mice using anti-human NEU1 and anti-human PPCA antibodies. Brown puncta depict compartmentalized/lysosomal NEU1 and PPCA. Scale bar: 50 µm. (E – N) Enzyme activity levels measured in the liver and spleen of WT, *Neu1^−/−^* and scAAV-N/P treated mice: NEU1 (E and F), CA (G and H), β-Gal (I and J), β-Hex (K and L) and α-Man (M and N) (*n* = 6). Values are expressed as mean ± SD. Statistical analysis was performed using the One-Way ANOVA; \**p* < 0.05, \*\**p* < 0.01, \*\*\**p* < 0.001, \*\*\*\**p* < 0.0001. (O - R) Representative IHC images of the liver (O) and spleen (P) from WT, *Neu1^−/−^* and scAAV-N/P treated mice stained with anti-Lamp1 (O and P) and anti-CD68 antibodies (Q and R).

As seen in other organs, the liver of untreated *Neu1^−/−^* mice showed increased Lamp1+ immunostaining in the hepatocytes, and most prominently in cells surrounding the bile duct and hepatic artery (Figure 7O). In the spleen, intense Lamp1 immunostaining was observed in areas of the red pulp that contained the most affected cells (Figure 7P). After treatment, Lamp1 levels in both liver and spleen were comparable to those in the WT tissues (Figures 7O and 7P). We next tested whether the inflammatory marker CD68 (Figures 7Q and 7R) would also be corrected. Indeed, both liver and spleen from the scAAV-N/P injected mice had reduced number of CD68+ cells, which was indicative of diminished or absent inflammation (Figures 7Q and 7R).

### Prevention of generalized fibrosis scAAV-treated Neu1^−/−^ mice

As final assessment of the extent of correction of the disease phenotype in scAAV N/P injected mice, we also performed Masson’s Trichrome staining of the kidney, liver and spleen and compared it with the staining of these tissues in untreated *Neu1^−/−^* mice and WT controls. As described previously, tissue sections of the *Neu1^−/−^* kidney stained with Masson’s Trichrome showed the presence of numerous collagen+ cells and increased connective tissue surrounding the glomeruli (Figure S7C). A similar staining pattern was seen in sections of the *Neu1^−/−^* liver that revealed collagen+ cells surrounding the bile ducts and the hepatic artery, as well as prominent collagen staining in the areas of the connective tissue lining the central hepatic vein (Figure S7D). In the *Neu1^−/−^* spleen the area with increased collagen was the red pulp (Figure S7E). At three months post treatment, no fibrosis was detected in the kidney, liver and spleen that appeared morphologically identical to the WT controls (Figures S7C – S7E). These results confirmed the occurrence of a pathogenic, generalized fibrotic condition in the connective tissue of *Neu1^−/−^* mice that can be used as a parameter to assess therapeutic efficacy after treatment.

## Discussion

AAV-mediated therapy has emerged as a reliable *in vivo* gene transfer system for LSDs, given its efficacy of infection, targeting both dividing and non-dividing cells, broad tropism, and safety. Upon successful infection, the genome of rAAV vectors inefficiently integrates into the host DNA and remains mostly episomal, facilitating expression of the encoded gene by the host’s transcription and translation machineries. There are multiple serotypes of the virus with distinct tissue tropism, which have been further engineered to either target specific cell populations or ubiquitously infect multiple cell types providing a more systemic effect ^24^. In addition, the use of cell-specific promoters restricts the expression of the transgene to only certain tissues or cells. The AAV serotype 2 was the first to be vectorized and several pseudo-types have been generated with cell- or tissue-specific tropism ^25,26^. These vectors achieve stable transgene expression, and have been extensively employed in pre-clinical studies on various disease models as well as in clinical trials^27^ ^28–45^ Several AAV-mediated gene therapy clinical trials, which have been approved by both the Food and Drug Administration and the European Medicines Agency,^24^ are currently in place for different LSDs, including Gaucher disease, GM1- and GM2-gangliosidoses, Krabbe disease, several mucopolysaccharidoses (I, II, IIIA, B, C, D, VII), metachromatic leukodystrophy, Pompe disease, Batten disease (neuronal ceroid lipofuscinoses; CLN1 – 8), Fabry disease, and Danon disease ^20,46–50^.

Although long-term studies in patients are still ongoing, the so far overall positive results of AAV-mediated gene therapy in other LSDs have encouraged the use of this approach for the treatment of sialidosis type I and II, which currently has no available cure. A study using scAAV2/8-LP1-*PPCA* vector in *Neu1^−/−^*;*NEU1^V54M^*mice, a mouse model of sialidosis type I with residual mutant NEU1 activity, showed widespread expression of PPCA in various organs at 1 month post injection^17^. Treatment with PPCA resulted in increased residual NEU1 activity in most tissues and reduced vacuolization in the kidney; however, the PPCA protein was not expressed in the brain, most likely due to short-term treatment (4 weeks) and route of injection^17^.

In the current study, treatment of *Neu1^−/−^* mice with concurrent injections of recombinant scAAV2/8-CMV-*NEU1* and scAAV2/8-CMV-*CTSA,* was designed with the idea that the localization, stability and activity of therapeutic, human NEU1 would be enhanced by co-expression of the human PPCA rather than the endogenous murine enzyme ^1,2,51^. The positive therapeutic outcome of this study confirms the validity of this approach for the treatment of sialidosis: a one-time IV co-injection of the two recombinant vectors not only cleared the progressive edema and extensive lysosomal expansion and vacuolization in tissues, but also prevented sialyl-oligosacchariduria and excessive deposition of collagen in the heart, skeletal muscle, liver, kidney and spleen. Most importantly, we found a measurable copy number of both viruses in different brain regions of the treated mice, which are notoriously difficult to target due to the selectivity of the blood brain barrier. Although this did not translate into a clear increase in NEU1 enzyme activity, a few disease phenotypes appeared to be fully or partially corrected. Complete reversal of the CP epithelium abnormalities likely accounts for the normalization of lysosomal exocytosis measured in the CSF of treated mice. In addition, the decreased expression of the neuroinflammatory marker CD68 in the brain of AAV-injected mice suggests that microgliosis was partially corrected. However, treatment did not resolve or prevent the progressive deposition of β-amyloid that is a prominent neurodegenerative phenotype of the *Neu1^−/−^* mice, again likely due to the limited levels of enzyme expressed by neurons in the brain parenchyma through IV injection. From these rather encouraging results, we can cautiously predict that this approach may have the potential to ameliorate some of the neuropathological signs characteristic of sialidosis in patients, such as myoclonus, seizures, gait abnormality and cerebellar ataxia ^4,5,52^.

As it is the case for other LSD patients, treatment for sialidosis, particularly type I, should start as soon as the patient is diagnosed to achieve the maximum effect of preventing disease progression and the development of debilitating neurological manifestations. However, given their often mild clinical symptoms that could be mistaken for other disease conditions, type I patients are often misdiagnosed or diagnosed when some of their clinical signs are already apparent. Nevertheless, AAV-mediated gene therapy for this category of patients may still be beneficial even if started later in life. In these cases, however, therapeutic efficacy may depend on the type of mutations and severity of the symptoms at the time of treatment.

We have shown that *Neu1^−/−^* mice develop generalized fibrosis affecting multiple organs. In the muscle, this phenotype is associated with severe muscle atrophy, which could be the cause of the hypotonia reported in some patients with sialidosis^4,13,15^. In absence of NEU1, fibrosis is driven by excessive release of exosomes, loaded with profibrotic signaling molecules, by myofibroblasts, ultimately resulting in muscle degeneration, and the activation of an inflammatory response^12,13,15^. Combined injection of the two AAV vectors effectively normalizes the levels of Lamp1 in the connective tissue of muscle and other organs. Since increased Lamp1 levels correlate with excessive lysosomal exocytosis, we can infer that normalization of Lamp1 is accompanied by reduced lysosomal exocytosis in the connective tissue. Indeed, we have shown previously that correction of Neu1 activity in skeletal muscle of *Neu1^−/−^* mice prevents the activation of the TGFβ signaling pathway and excessive production and release of collagens ^13^. Therefore, we are confident that the overall reversal of the fibrotic disease seen here not only in muscle but also in heart, liver, kidney and spleen of *Neu1^−/−^* treated mice must be due at least in part to the decreased exocytic activity of fibroblasts and the reduction of inflammation^13^. This could be particularly relevant for patients with sialidosis type II who may develop a heart condition characterized by heart murmur ^6,53,54^, dilated ventricles and atria, decreased ventricular function ^6,54–60^ and abnormality of the mitral valves ^6,54,58,61^ that may lead to heart failure. The restoration of connective tissue morphology in the heart of *Neu1^−/−^* mice after treatment suggests that an AAV-mediated gene therapy approach may prevent heart involvement in some of the sialidosis patients. Furthermore, in the adult human population down-regulation of NEU1 may represent a risk factor for the initiation and/or progression of idiopathic forms of fibrosis, including idiopathic pulmonary fibrosis, and generalized fibrosis of the liver and kidney ^13^. In these cases, increasing the endogenous levels of NEU1 may be beneficial to prevent or stop the progression of the disease, although more studies are needed to demonstrate the involvement of NEU1 in human forms of fibrosis.

We have previously shown that a recombinant NEU1 enzyme used for ERT in *Neu1^−/−^* mice elicits a severe immune response^16^, which would raise concerns about the development of adverse immunologic reactions in patients. However, this therapeutic approach would still be valid for Type I patients with residual enzyme activity. In this study, no immune response was mounted against NEU1 and PPCA, albeit antibodies were detected against the viral vector. These results are in line with recently reported data on a long-term safety and efficacy study, using a liver-specific scAAV2/8-LP1-*PPCA* vector to treat the mouse model of galactosialidosis (*Ppca^−/−^*). Although no neutralizing antibodies against the PPCA protein were detected 12 months after injection, antibodies against viral capsid were measured; however, this immune response did not affect protein expression levels or the efficacy of treatment^62^

Wild-type AAV infections are common in humans starting from the first year of life, although they are not associated with any disease^63–65^. These infections cause a humoral response against the virus that results in neutralizing antibodies, which can reduce the efficacy of the rAAV serotypes used for treatment, ultimately lowering the number of patients eligible for treatment ^64,66^. In addition, neutralizing antibodies generated post injection with long-term persistence can prevent re-administration of the same rAAV or other rAAV vectors when needed ^64^. To overcome this complication, drugs that block humoral or cellular immune response, have been used in conjunction with systemically delivered gene therapies ^64,66^. Many of these immunomodulatory or suppressive drugs are currently in preclinical trials, clinical trials (as combination therapy), or approved ^66^. These drugs, alone or in combination, have been shown to reduce the number of neutralizing antibodies, block both T or B cell responses, and prevent the formation of anti-capsid and anti-transgene antibodies, thus allowing rAAV readministration when experiencing an expression loss ^66^.

The success of these preclinical studies supports the development of this therapeutic approach for the treatment of sialidosis. In addition, given the beneficial effects of repurposed drugs for ameliorating some of the clinical signs in type I sialidosis patients, a controlled diet and medication regimen should be taken into consideration in combination with target therapy, including gene therapy, for this disease.

## Materials and Methods Animals

Animals were housed in a fully AAALAC (Assessment and Accreditation of Laboratory Animal Care) accredited animal facility with controlled temperature (22 °C), humidity, and lighting (alternating 12 h light/dark cycles). Food and water were provided ad libitum. All procedures in mice were performed according to animal protocols approved by the St. Jude Children’s Research Hospital Institutional Animal Care and Use Committee and National Institutes of Health guidelines (#235-100636-02/23; date of approval 02-22-2023). WT and *Neu1^−/−^* mice aged 1-, 2-, and 4-months (FVB/NJ background)^7^ were included in this study. One-month-old mice were injected with a single combined dose of scAAV2/8-CMV-*NEU1* (2×10^12^ vg/mouse) and scAAV2/8-CMV-*CTSA* (1×10^12^ vg/mouse) (scAAV2/8-N/P) and sacrificed at 4 months of age.

### Production and titration of the scAAV2/8-CMV-*NEU1* and scAAV2/8-CMV *-CTSA* vectors

The production and titration of the scAAV2/8-CMV-*NEU1* and scAAV2/8-CMV-*CTSA* constructs has been described previously^62^. The proteins are under the control of a constitutive cytomegalovirus early enhancer promotor that drives the expression of the 1248 base pairs human *NEU1* and 1443 base pairs human *CTSA* cDNAs, respectively. The scAAV vector particles were made using a good manufacturing practice (GMP) process, comparable transient-transfection procedure in the Children’s GMP, LLC facility on the St. Jude campus, as previously described for the hemophilia B vector^67^.

### Biodistribution

Total genomic DNA was extracted from the cerebellum, cortex, brain stem, hippocampus, liver, lung, heart, kidney, spleen, ovary, diaphragm, jejunum, colon, peritoneum, inguinal lymph nodes and by using the DNeasy Blood and Tissue Kit (Qiagen, Germantown, MD), following the manufacturer’s guidelines. RT-qPCR was performed using SsoAdvanced universal SYBR green supermix (Bio-Rad, Hercules, CA), 100 ng or 250 ng genomic DNA, 10 µM primers (CMV promotor: 5’– CGG TGG GAG GTC TAT ATA A –3’; PPCA: 5’– GCG GGC TCG GAT CAT GGT GGA AT –3’; NEU1: 5’– GTC GCT CCC CAG TCA TCT CTC CC –3’) and RNAse free water in a 25 µL reaction volume on a CFX96 opus 96 Real-Time PCR system (Bio-Rad, Hercules, CA). The copy number was calculated using the molecular weight of CMV-PPCA or CMV-NEU1 plasmids. The standard curve was generated for quantification of the viral genome. PPCA standard curve: 1.79×10^8^, 8.79×10^7^, 1.76×10^7^, 8.79×10^6^, 1.76×10^6^, 8.79×10^5^, 1.76×10^5^, 8.79×10^4^, 1.79×10^4^, 8.79×10^3^, 1.79×10^3^ copies. NEU1 standard curve: 1.84×10^8^, 9.19×10^7^, 1.84×10^7^, 9.19×10^6^, 1.84×10^6^, 9.19×10^5^, 1.84×10^5^, 9.19×10^4^, 1.84×10^4^, 9.19×10^3^, 1.84×10^3^copies. All samples were tested in triplicate and a negative control (H_2_O) was included to monitor sample contamination.

### Detection of antibodies against AAV2/8 capsid and human NEU1 and PPCA transgene products

Antibody levels were tested in the sera of WT, *Neu1^−/−^* mice, and treated *Neu1^−/−^* mice via an enzyme-linked immunosorbent assay (ELISA). Serum collected at 3 months post injection was used for the ELISA. Briefly, either disrupted or intact scAAV2/8–CMV–*NEU1* and scAAV2/8– CMV–*CTSA* virus or purified NEU1 (Novus Biologicals, Centennial, CO) and PPCA proteins was diluted to 1 mg/mL in DPBS at 50 μL per well in a Nunc MaxiSorp ELISA plate (Thermo Scientific, Waltham, MA). When purified virus was used, virus concentration was estimated using NanoDrop lite Spectrophotometer (Thermo Scientific, Waltham, MA), prior to disruption. The virus stock was incubated in disruption buffer containing 5 mM Tris (pH 7.8), 60 mM KCl, and 0.05% Triton X-100 for 15 min at RT. Virus was then diluted in DPBS to 1 mg/mL. A human specific NEU1 or PPCA antibody was used as positive control for the ELISA against the two proteins and an anti-AAV8 antibody (Progen, Wayne, PA) was used as a positive control against the capsid. Plates were incubated at 4 °C overnight to allow the protein/virus to bind. Following incubation, the plates were washed twice with DPBS/ 0.05% Tween 20 (washing buffer) and blocked with 10% FBS in DPBS (blocking buffer) for 30 min at RT. Plates were washed twice with washing buffer, and diluted samples in triplicate were placed in the wells for 2h. Plates were then washed twice with washing buffer, and goat anti-mouse immunoglobulin G-horseradish peroxidase (IgG-HRP) (Southern Biotechnology, Birmingham, AL) was used as a secondary antibody diluted 1:5,000 in blocking buffer. Plates were incubated for 1 h at RT and washed three times with washing buffer, and TMB 2-Component Microwell Peroxidase Substrate (SeraCare Life Sciences, Gaithersburg, MD) was added to each well. Plates were allowed to develop for 15 min, and the reaction was stopped by adding an equal volume of 1 M phosphoric acid. Plates were then read in a Molecular Devices Spectramax M5 (Molecular Devices, San Jose, CA) at an OD of 450. Triplicates were averaged and plotted.

### Immunohistochemical analyses

For IHC analysis, tissues were fixed in 10% buffered formalin and embedded in paraffin. The sections were cut (6 µm) and deparaffinized. For NEU1, PPCA, and CD68 staining, antigen retrieval was performed by the pressure cooker method using citrate buffer [0.1 M citric acid and 0.1 M sodium citrated (pH 6.0) containing 0.05% Tween-20]. After sections were blocked with 2.5% Normal Horse Serum (Vector Laboratories, Newark, CA), they were incubated overnight at RT with anti-human NEU1 (in-house), anti-human PPCA (in-house), or anti-mouse CD68 (Cell Signaling Technologies, Danvers, MA) antibodies. Sections were rinsed in PBS and subsequently incubated with biotinylated secondary goat anti-rabbit antibody (Jackson ImmunoResearch Laboratory, West Grove, PA) for 1 h. Endogenous peroxidase was quenched by incubation the sections with 3% hydrogen peroxidase for 15 min. Sections were then developed using the ImmPRESS® HRP Horse Anti-Rabbit IgG Polymer Detection Kit (Vector Laboratories, Newark, CA) and counterstained with hematoxylin.

For Lamp1 staining, antigen retrieval was completed in the microwave using citrate buffer [0.1 M citric acid and 0.1 M sodium citrated (pH 6.0) containing 0.05% Tween-20]. After sections were blocked with PBS containing 0.1% bovine serum albumin (BSA), 10% Normal Goat Serum and 0.5% Tween-20, they were incubated overnight at RT with anti-mouse Lamp1 (Cell Signaling Technologies, Danvers, MA) antibody. Sections were rinsed in PBS and subsequently incubated with biotinylated secondary goat anti-rabbit antibody (Jackson ImmunoResearch Laboratory, West Grove, PA) for 1 h. Endogenous peroxidase was quenched by incubation the sections with 3% hydrogen peroxidase for 15 min. Sections were then developed using the ImmPRESS® HRP Horse Anti-Rabbit IgG Polymer Detection Kit and counterstained with hematoxylin.

For APP staining the Mouse on Mouse (M.O.M.) ImmPRESS HRP Polymer Kit (Vector Laboratories Newark, CA) was used. Antigen retrieval was performed by the pressure cooker method using citrate buffer [0.1 M citric acid and 0.1 M sodium citrated (pH 6.0) containing 0.05% Tween-20]. Endogenous peroxidase was quenched by incubation the sections with 3% hydrogen peroxidase in methanol for 15 min. Slides were washed twice in PBS for 5 min and incubated for 1 h in the working solution of prepared M.O.M. mouse IgG blocking reagent. Sections were washed twice in PBS for 5 min and incubated for 5 min in M.O.M. Normal Horse Serum, 2.5%. Sections were incubated overnight at RT with mouse anti-APP antibody for 30 min. The sections were washed twice in PBS and incubated for 10 min with M.O.M. ImmPRESS Reagent and washed again twice with PBS. Antibody detection was performed using the ABC Kit (Vector Laboratories, Newark, CA) and diaminobenzidine substrate (Invitrogen, Carlsbad, CA) and counterstained with hematoxylin.

For histopathological examination, sections were cut (6 µm) and stained with a standard H&E method.

### Enzyme assays

The tissues were homogenized in 10x w/v ddH_2_O and, when necessary, further diluted in Enzyme Dilution Buffer (50 mM Na-Acetate pH 5.0, 100 mM NaCl, 1 mg/ml BSA (w/v), 0.02% Na-azide (w/v)). Enzyme activities were measured with the appropriate fluorometric substrates as previously described ^18,68,69^. Neuraminidase 1 (Neu1) activity was measured against synthetic 4-methylumbelliferyl-α-D-N-acetylneuraminic acid (Sigma-Aldrich, St. Louis, MO) at pH4.3, beta-galactosidase (β-Gal) activity against 4-methylumbelliferryl-β-D-Galactopyranoside (Sigma-Aldrich, St. Louis, MO), beta-hexosaminidase (β-Hex) activity against 4MU-N-acetyl-β-D-glucosaminide (Sigma-Aldrich, St. Louis, MO), alpha-mannosidase (α-Man) activity against 4MU-α-D-mannopyranoside (Sigma-Aldrich, St. Louis, MO), and cathepsin A (CA) activity against the dipeptide Z-Phe-Ala (Sigma-Aldrich, St. Louis, MO). All reactions were performed at 37 °C for 1 h and stopped with carbonate stop buffer (0.5 M Na_2_CO_3_ with the pH set to 10.7 by adding 0.5 M NaHCO_3_) for Neu1, β-Gal, β-Hex, and α-Man activities and the CA activity was stopped at 100 °C for 5 min and 10 µL reaction mixture was read in 250 µL 50 mM Na-carbonate stop buffer pH9.5 containing 500 µL o-phtaldialdehyde (10 mg/mL) and 500 µL 2-mercaptoethanol (5 mL/mL) per 30 mL. All fluorescence was measured at λ_ex_=355/λ_em_=460 and interpolated to a standard curve adjusted for dilution, length of incubation, and normalized to protein concentration.

### Masson Trichrome Staining

Masson’s trichrome staining was performed according to the manufacturer’s protocol (Polysciences Inc., Warrington, PA). Tissues were fixed in 10% buffered formalin and embedded in paraffin. were cut (6 µm), deparaffinized, and fixed in Mordant in Bouin’s solution for 1 h at 60 °C. Sections were stained sequentially at room temperature with Weigert’s iron hematoxylin (10 min), Biebrich scarlet acid fuchsin (5 min), phosphotungstic /phosphomolybdic acid (10 min), and aniline blue (5 min). The sections were washed, dehydrated, and mounted with a xylene-based mounting medium.

### Sialic Acid assay

Urine was collected from individual control animals (WT and *Neu1^−/−^*) at 1-, 2- and 4-months of age and from 2- and 4-month-old mice treated with scAAV-N/P. Urine samples were further diluted (1:5) in PBS when necessary. The total sialyloligosaccharide content in mouse urine was determined by the release of bound sialic acid using an Enzychrome Sialic Acid Assay Kit from BioAssay Systems (Bioassay Systems, Hayward, CA). Total sialic acid was measured following acid hydrolysis for 1 h at 80 °C. After the addition of neutralization buffer, samples were cooled at RT and incubated with an enzymatic master mix to utilize an enzyme-coupled reaction to oxidize any liberated sialic acid. A standard curve was generated from a 10 mM sialic acid stock solution that was serially diluted in ultrapure water. The fluorescence was read at λ_ex_=535nm, λ_em_=595nm and the data were interpolated to the standard curve to yield a concentration value in µM (nmol/mL).

### Vacuolization area measurements

To measure the extent of vacuolization in kidney sections, 1-month, and 4-months old WT and *Neu1^−/−^* mice and 4-months old mice treated with scAAV-N/P were stained for H&E (*n* = 8). A macro was developed in ImageJ to analyze six random areas per section of each mouse. A detection threshold was set for each genotype to analyze the total area occupied by vacuoles and batched to all images using the same parameters.

### Statistical analysis

Statistical analyses were performed using GraphPad Prism 9 (GraphPad Software, San Diego, CA). Significant differences between study groups for all quantitative outcome measures were established using One-way ANOVA or the Brown-Forsythe and Welch ANOVA (Figure 2G – 2K). Significance was defined as *p* < 0.05. The data are presented as mean ± SD.

### Data availability

All relevant data supporting the key findings of this study are available within the article and its supplemental information files or from the corresponding author upon reasonable request.

## Acknowledgements

A.d’.A. holds the Jewelers for Children Endowed Chair in Genetics and Gene Therapy. This work was funded by NIH grants R01GM104981; R44NS084565; 1RF1NS123174; the Assisi Foundation of Memphis and the American Lebanese Syrian Associated Charities (ALSAC).

## Author Contributions

Conceptualization, D.v.d.V., Y.C. and A.d’A.; Methodology, D.v.d.V., L.E.F., H.H., J.A.W., S.A.B., and A.d’A.; Validation, D.v.d.V. and A.d’A.; Formal Analysis, D.v.d.V. and J.A.W.; Investigation, D.v.d.V., L.E.F., H.H., J.A.W. and S.A.B.; Resources, D.v.d.V., L.E.F., H.H., J.A.W., S.A.B., E.G. and A.d’A.; Software, J.A.W.; Data Curation, D.v.d.V., L.E.F., J.A.W. and A.d’A.; Writing – Original Draft Preparation, D.v.d.V.; Writing – Review & Editing, A.d’A D.v.d.V.; Visualization, D.v.d.V. and A.d’A.; Supervision, A.d’A.; Project Administration, A.d’A.; Funding Acquisition, A.d’A. All authors have read and agreed to the published version of the manuscript.

