## Supplemental material for "AAV-mediated gene therapy for Sialidosis"

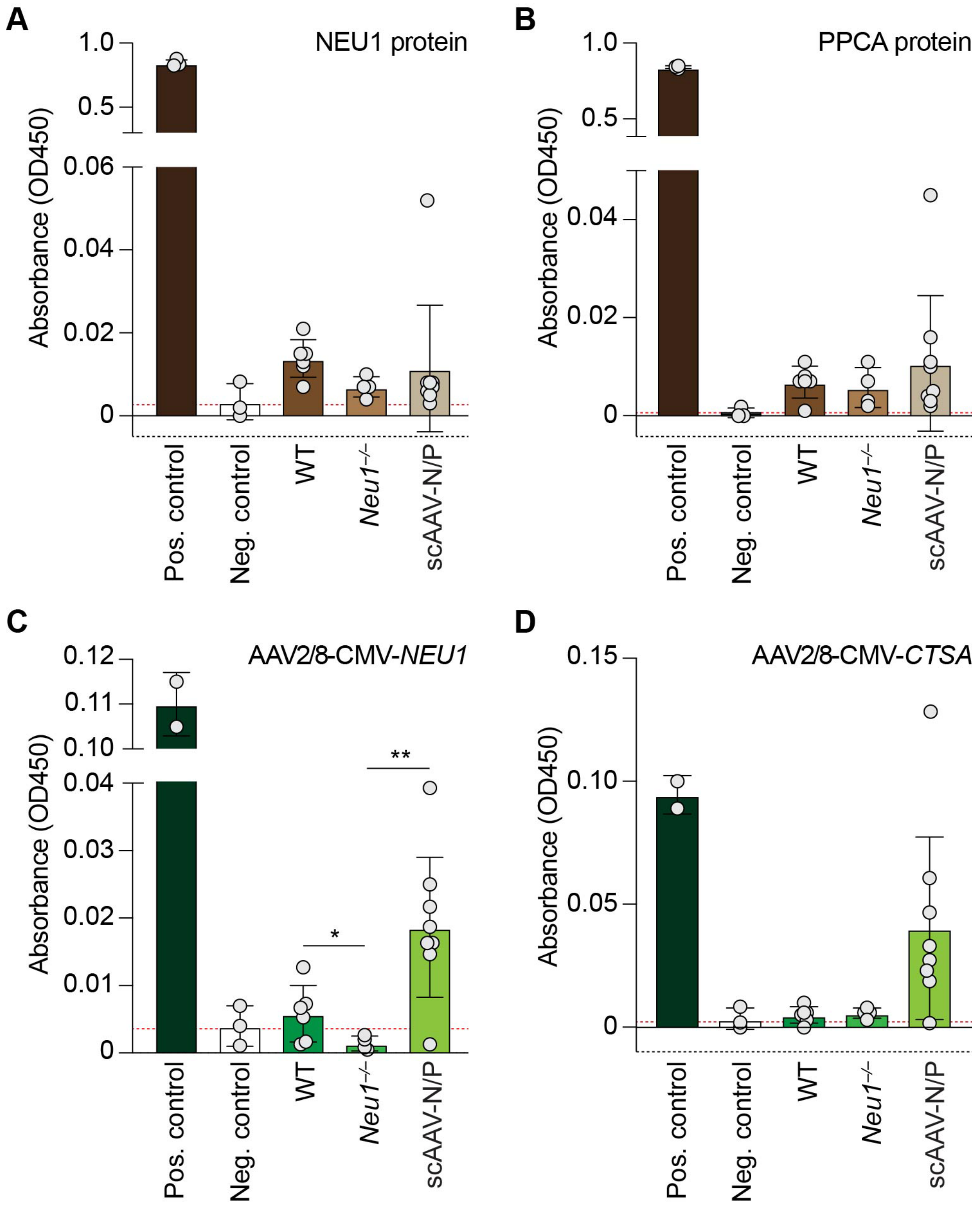


**Figure S1. Antibody of measurements in sera of scAAV-N/P treated mice**. (A – D) Levels of neutralizing antibodies against the NEU1 (A) and PPCA (B) transgene products and AAV-2/8 capsid (C and D) were assessed by ELISA. Anti-NEU1, anti-PPCA and anti-AAV8 antibodies were used as positive controls. Sera samples were diluted 1:50 (capsid) and 1:100 (protein). WT $n$ = 6, *Neu1*$^{-/-}$ $n$ = 4 and scAAV-N/P $n$ = 8. Values are expressed as mean  SD. Statistical analyses were performed using One-way ANOVA; *$p < 0.05$, **$p < 0.01$.

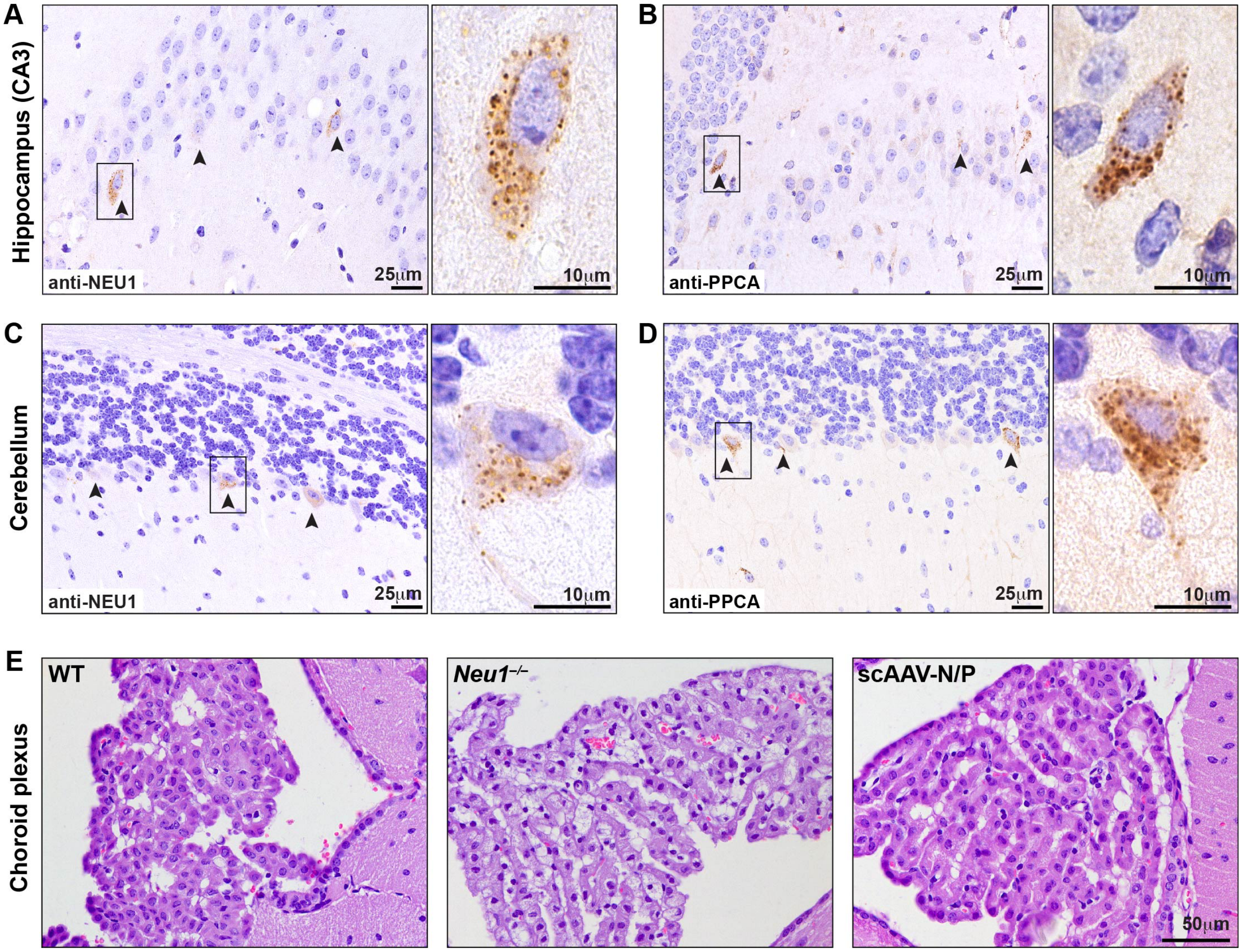


**Figure S2. Restoration of choroid plexus morphology in scAAV-N/P treated mice.** (A – D) NEU1 and PPCA protein levels in some neurons within the hippocampal CA3 region (A and B) and Purkinje cells in the cerebellum (C and D). Scale bar: 25 □m. Right panels are zoomed in images of boxed areas. Scale bar: 10 □m. Brown puncta depict compartmentalized/lysosomal NEU1 and PPCA. (E) Representative images of H&E-stained sections of CP from WT, *Neu1*$^{-/-}$ and scAAV-N/P treated mice. Scale bar: 50 µm.

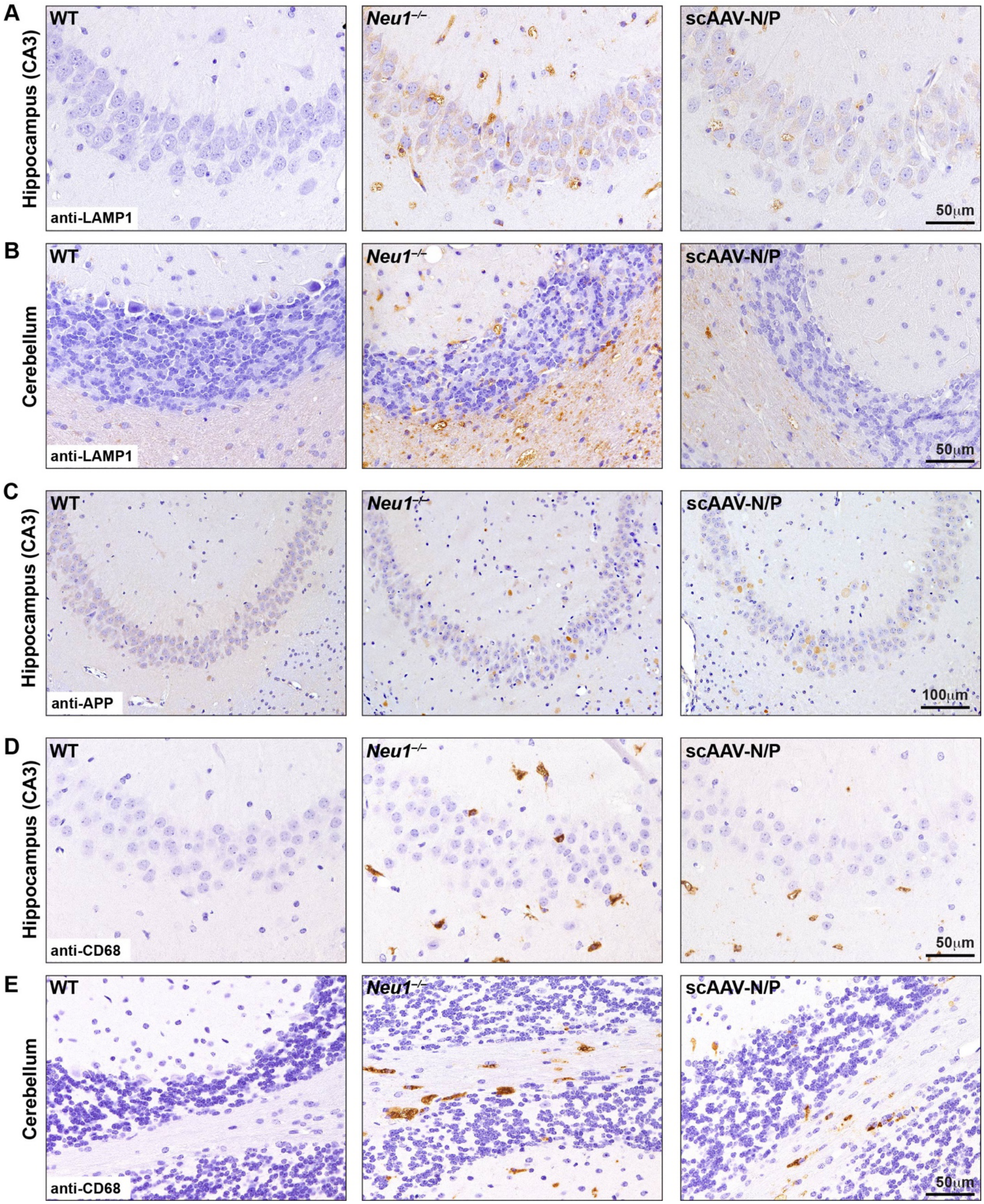

A
Hippocampus (CA3)
WT
Neu1−/−
scAAV-N/P
anti-LAMP1
50µm
B
Cerebellum
WT
Neu1−/−
scAAV-N/P
anti-LAMP1
50µm
C
Hippocampus (CA3)
WT
Neu1−/−
scAAV-N/P
anti-APP
100µm
D
Hippocampus (CA3)
WT
Neu1−/−
scAAV-N/P
anti-CD68
50µm
E
Cerebellum
WT
Neu1−/−
scAAV-N/P
anti-CD68
50µm

**Figure S3. Reduced Lamp1 and CD68 expression in brain regions from the scAAV-N/P treated mice.** (A – B) Representative IHC images of the CA3 region of the hippocampus (A) and cerebellum (B) from WT, *Neu1*$^{-/-}$ and scAAV-N/P treated mice using anti-Lamp1 antibody. Scale bar: 50 μm. (C) Representative IHC images of the CA3 region of the hippocampus from WT, *Neu1*$^{-/-}$ and scAAV-N/P treated mice using anti-APP antibody. Scale bar: 50 μm (C – D) Representative IHC images of the CA3 region of the hippocampus (C) and cerebellum (D) from WT, *Neu1*$^{-/-}$ and scAAV-N/P treated mice using anti-CD68 antibody. Scale bar: 50 μm

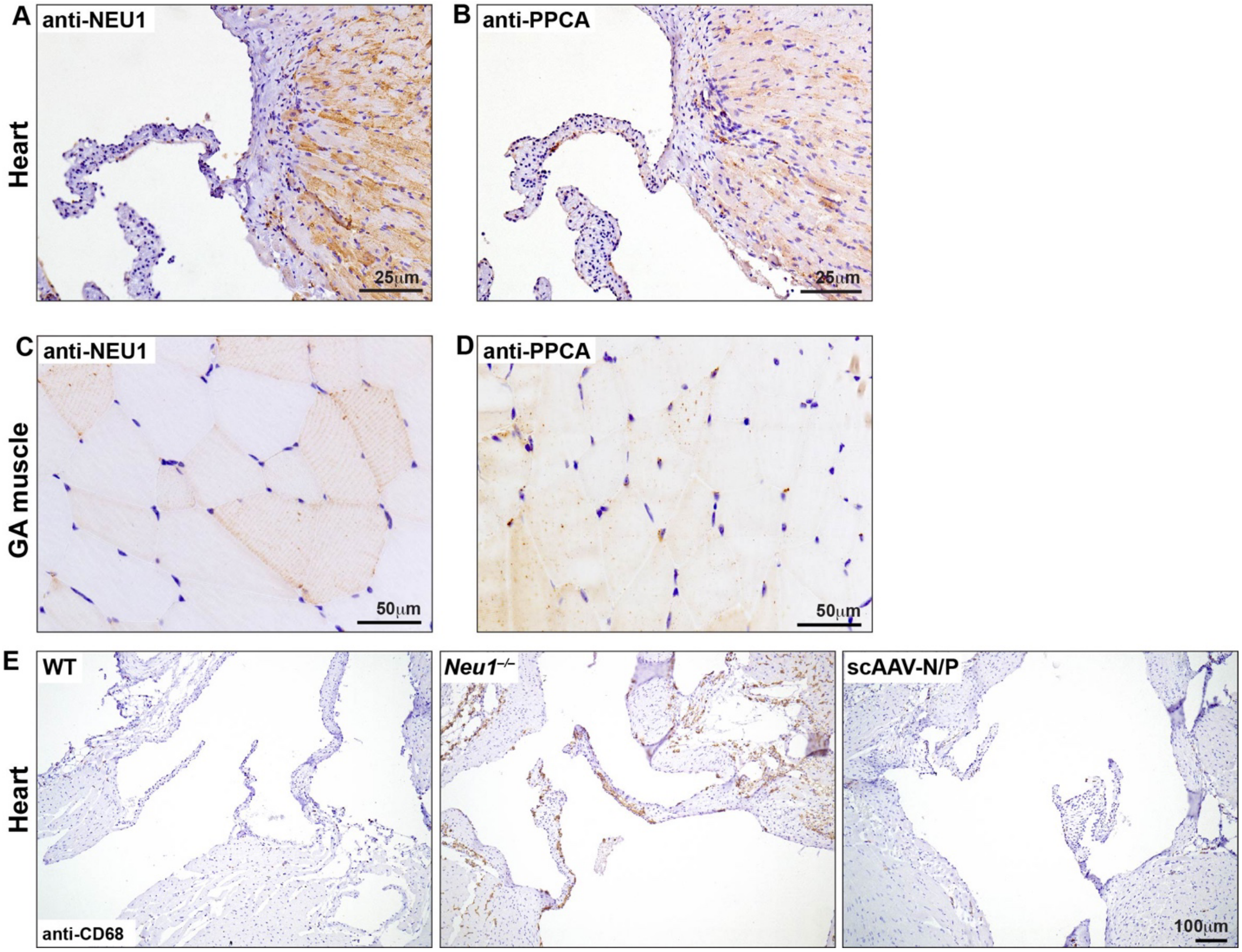


**Figure S4. NEU1 and PPCA levels in heart and skeletal muscle of treated mice.** (A – D) Representative IHC images of the heart valves (A and B) and coronal sections of the skeletal muscle (C and D) using human anti-NEU1 and anti-PPCA antibodies. Scale bars: 25 μm and 50 μm. (E) Representative IHC images of the heart valves anti-Lamp1 antibody. Scale bar: 100 μm.

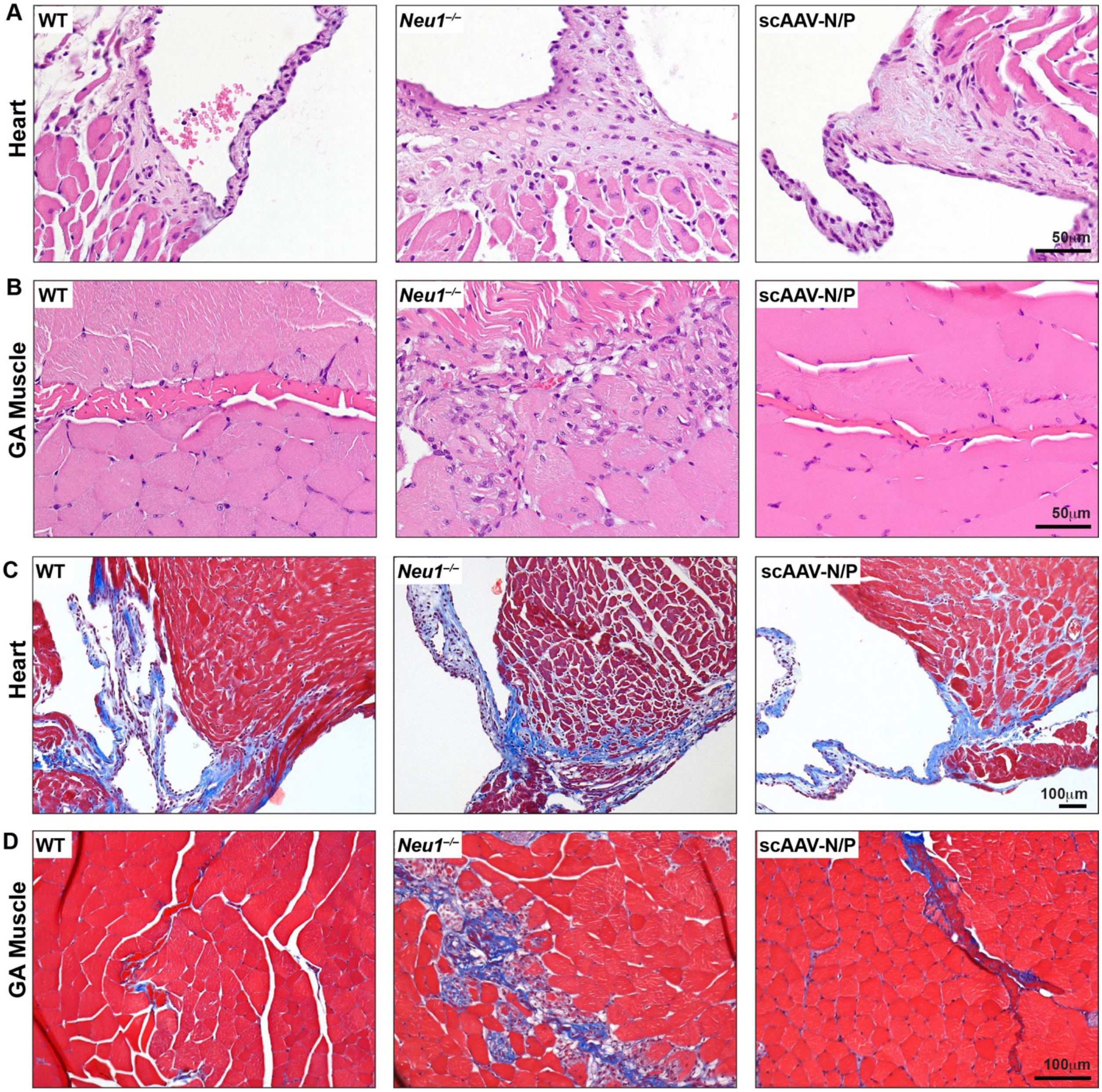


**Figure S5. Correction of fibrotic disease in heart and skeletal muscle of scAAV-N/P treated mice**. (A and B) Representative H&E images of the heart valves (A) and GA skeletal muscle (B) showing normalized morphology comparable to the WT control. Scale bar: 50 µm. (C and D) Masson's trichrome stained heart (C) and GA skeletal muscle (D) sections showing complete resolution of muscle fibrosis. Scale bars: 50 µm and 100 µm.

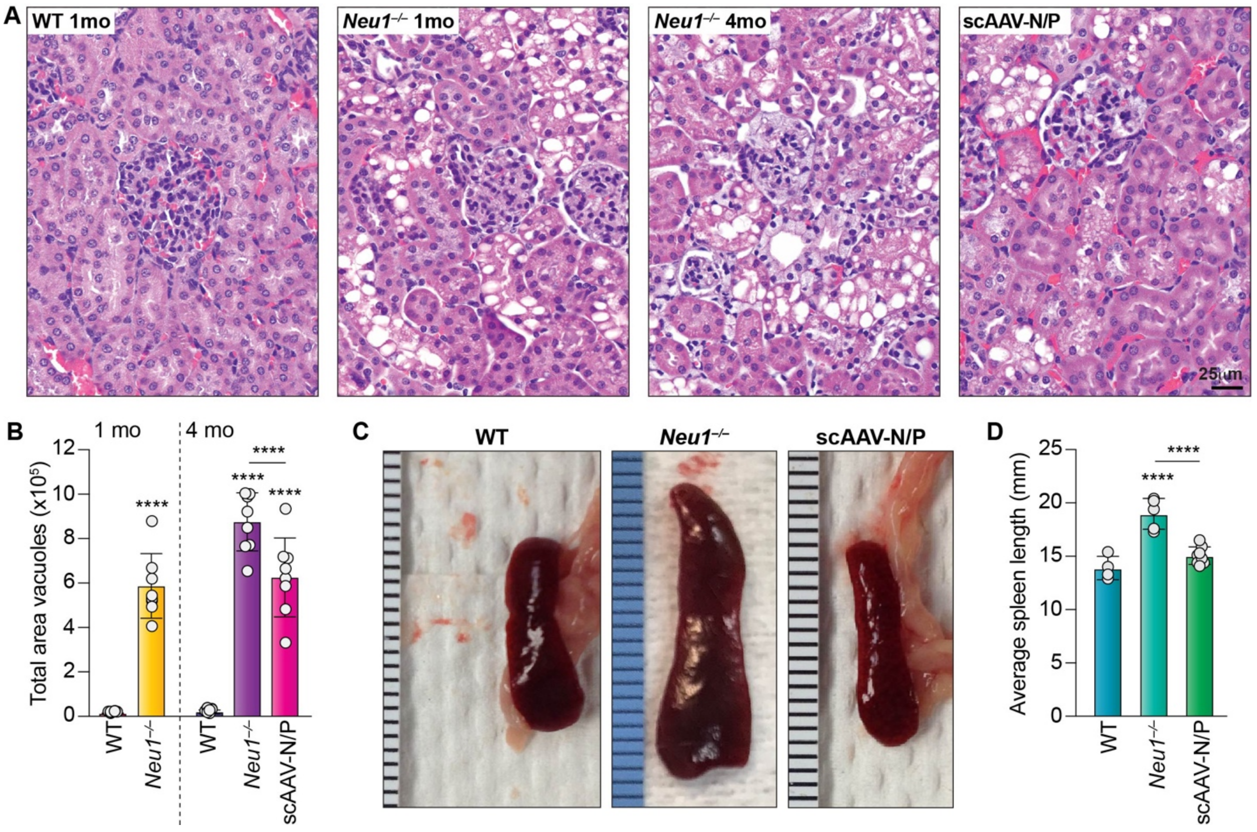


**Figure S6. Reduction of vacuolization in the kidney and prevention of splenomegaly in treated mice.** (A) Representative H&E images of WT (1 mo), *Neu1*$^{–/–}$ (1 mo), *Neu1*$^{–/–}$ (4 mo) and *Neu1*$^{–/–}$-scAAV-N/P (4 mo) kidneys. Scale bar: 25 µm. (B) Quantification of the vacuolated areas in the kidneys before (at 1-month) and after treatment (at 4-months) ($n$ = 8). Values are expressed as mean □ SD. Statistical analysis was performed using the One-Way ANOVA; ****$p$ < 0.0001. (C and D) Spleen size measured in WT, *Neu1*$^{–/–}$ and scAAV-N/P treated mice (C) and statistical assessment of spleen size from multiple treated and untreated mice and controls (D). WT; $n$ = 4, *Neu1*$^{–/–}$; $n$ = 5 and scAAV-N/P treated; $n$ = 8. Values are expressed as mean □ SD. Statistical analysis was performed using the One-Way ANOVA; ****$p$ < 0.0001.

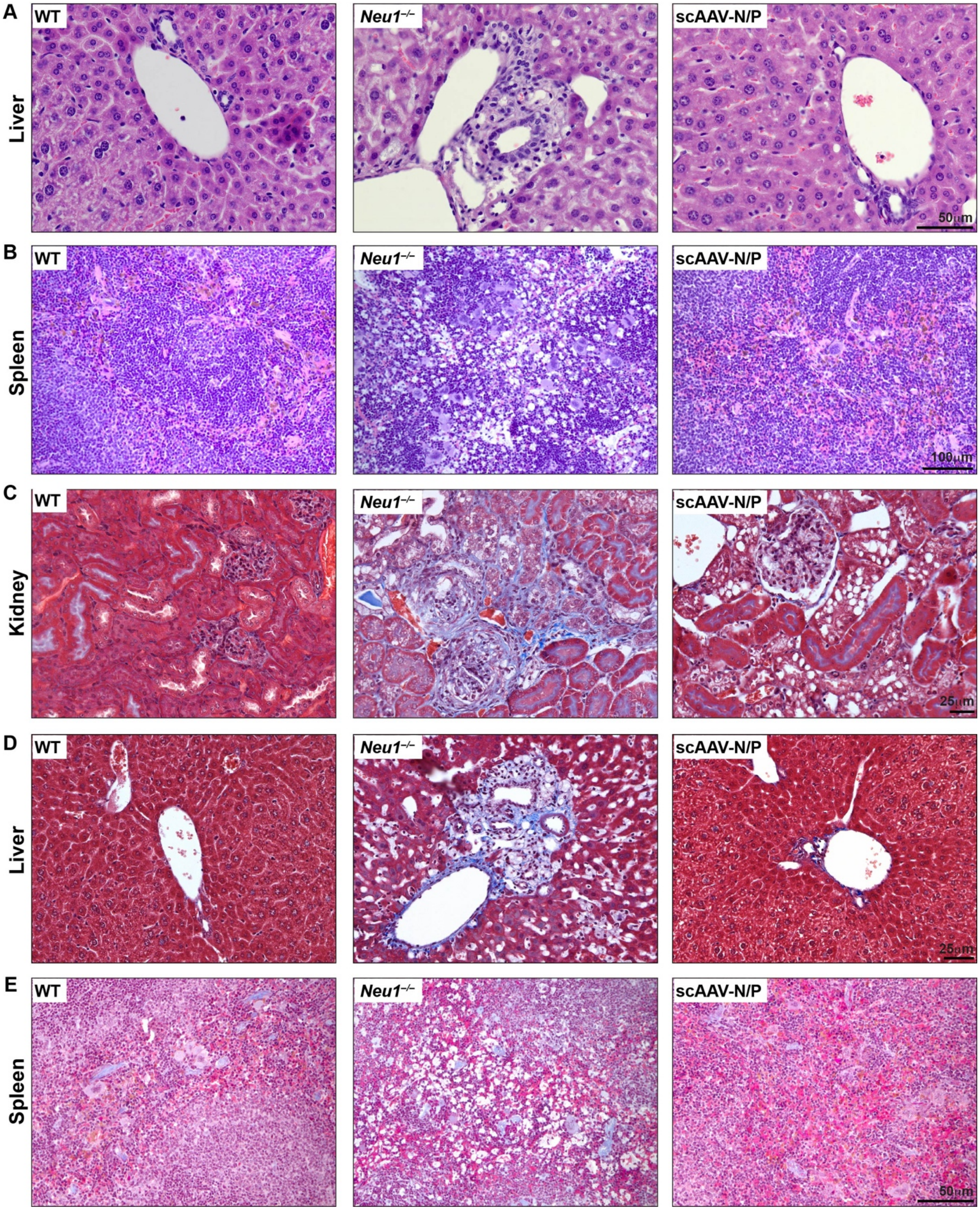
A
B
C
D
E
Liver
Spleen
Kidney
Liver
Spleen
WT
Neu1–/–
scAAV-N/P
50μm
100μm
25μm
25μm
50μm

**Figure S7. Restored morphology of the liver and spleen and prevention of collagen deposition in kidney, liver and spleen of treated mice**. (A) Representative images of H&E-stained liver sections from WT, *Neu1*$^{-/-}$ and scAAV-N/P treated. Scale bars: 50 µm. (B) Representative images of H&E-stained spleen sections from WT, *Neu1*$^{-/-}$ and scAAV-N/P treated mice. Scale bar: 100 µm. (C - F) Masson's Trichrome stained sections of the kidney (C), liver (D) and spleen (E) from WT, *Neu1*$^{-/-}$, and scAAV-N/P mice. Scale bars: 25 µm and 50 µm.
